# Lipid droplets control mitogenic lipid mediator production in human cancer cells

**DOI:** 10.1101/2021.11.25.470010

**Authors:** Eva Jarc Jovičić, Anja Pucer Janež, Thomas O. Eichmann, Špela Koren, Vesna Brglez, Paul M. Jordan, Jana Gerstmeier, Duško Lainšček, Anja Golob-Urbanc, Roman Jerala, Gérard Lambeau, Oliver Werz, Robert Zimmermann, Toni Petan

## Abstract

Polyunsaturated fatty acids (PUFAs) are components of membrane phospholipids and precursors of bioactive lipid mediators. Here, we investigated the crosstalk of three pathways providing PUFAs for lipid mediator production: (i) secreted group X phospholipase A_2_ (GX sPLA_2_) and (ii) cytosolic group IVA PLA_2_ (cPLA_2_α), which both mobilize PUFAs from phospholipids, and (iii) adipose triglyceride lipase (ATGL), which breaks down triacylglycerols (TAGs) stored in lipid droplets (LDs). Combining lipidomic and functional analyses, we demonstrate that lipid mediator production depends on TAG turnover. GX sPLA_2_ directs PUFAs into TAGs and ATGL is required for their entry into lipid mediator biosynthetic pathways. ATGL also promotes the incorporation of LD-derived PUFAs into phospholipids representing substrates for cPLA_2_α. Additionally, inhibition of TAG synthesis mediated by acyl-CoA:diacylglycerol acyltransferase 1 (DGAT1) reduces the levels of mitogenic lipid signals and compromises tumour growth. This study expands the paradigm of PLA_2_-driven lipid mediator signalling and identifies LDs as central lipid mediator production hubs.

## Introduction

Fatty acids (FAs) are universal cellular energy sources and membrane building blocks. FAs are also involved in signalling pathways that control cell growth, inflammation and tumourigenesis (Röhrig & Schulze, 2016). The ensemble of signalling FAs is vastly expanded by the oxygenation of polyunsaturated FAs (PUFAs) by cyclooxygenase (COX), lipoxygenase (LOXs) and CYP450 monooxygenase enzymes into several families of bioactive lipid mediators, including the eicosanoids (Dennis & Norris, 2015). These short-lived autocrine and paracrine signalling molecules are released from cells to collectively modulate various processes in their microenvironment, e.g., to orchestrate a shift into pro-inflammatory or anti-inflammatory states. Pathways that control the availability of different PUFAs for oxygenation determine the types of lipid mediator species produced and the dynamics of their production (Wang & DuBois, 2010; Greene *et al*, 2011). Our current understanding of the control of PUFA supply for lipid mediator production is limited, particularly as this is intrinsically dependent on complex (PU)FA metabolism, including PUFA biosynthesis, uptake, storage, breakdown, lipid remodelling and trafficking (Pérez-Chacón *et al*, 2009; Astudillo *et al*, 2019; Jarc & Petan, 2020).

The canonical pathway that supplies arachidonic acid (AA) for eicosanoid production depends on the group IVA cytosolic phospholipase A_2_ (cPLA_2_α), whose role in stimulus-induced eicosanoid production has been demonstrated in various pathophysiological settings (Bonventre *et al*, 2004; Shimizu, 2009; Murakami *et al*, 2011; Leslie, 2015). Upon cell activation, cPLA_2_α binds to perinuclear membranes of the ER and Golgi complex and selectively hydrolyses phospholipids containing AA at the *sn-2* position (Hayashi *et al*, 2021). Numerous other members of the PLA_2_ superfamily promote lipid mediator production, either through activation of cPLA_2_α or by acting independently on their respective phospholipid pools, thereby releasing not only AA but also other PUFAs (Saiga *et al*,2005; Duchez *et al*, 2019; Astudillo *et al*, 2019). In particular, several secreted PLA_2_s (sPLA_2_s) have been implicated in lipid mediator production (Murase *et al*, 2016; Sato *et al*, 2020). The group X sPLA_2_ is the most potent among mammalian sPLA_2_s at hydrolysing the phosphatidylcholine (PC)-rich plasma membrane, lipoproteins and extracellular vesicles (Bezzine *et al*, 2002; Mounier *et al*, 2004; Jarc *et al*, 2018; Kudo *et al*, 2022). It releases various unsaturated FAs, including ω-3 and ω-6 PUFAs, and it is involved in inflammation, immunity and cancer (Surrel *et al*, 2009; Murase *et al*, 2016; Schewe *et al*, 2016; Pucer *et al*, 2013; Ogden *et al*, 2020; Kudo *et al*, 2022).

Recent studies suggest that besides membrane phospholipids, other cellular lipid pools, including neutral lipids stored in lipid droplets (LDs) or derived from lipoprotein uptake, are also sources of PUFAs for lipid mediator production (Dichlberger *et al*, 2014; Schlager *et al*, 2015, 2017; Jarc & Petan, 2020). LDs are specialized organelles that store FAs and other lipids in their esterified forms, primarily as triglycerides (TAGs) and sterol esters (Welte & Gould, 2017; Olzmann & Carvalho, 2019). A hallmark role of LDs is taking up excess FAs to prevent lipotoxicity and fine-tune FA release via lipolysis to match various cellular demands (Listenberger *et al*, 2003; Rambold *et al*, 2015; Nguyen *et al*, 2017; Henne *et al*, 2018; Petan, 2020). Adipose triglyceride lipase (ATGL), the major mammalian TAG lipase, provides FAs for mitochondrial energy production, but it also controls FA-induced signalling pathways that coordinate metabolism and inflammation (Zimmermann *et al*, 2004; Haemmerle *et al*, 2011; Schweiger *et al*, 2017; Zechner *et al*, 2017). In immune and cancer cells, LDs have long been implicated in AA storage, trafficking and eicosanoid production (Dvorak *et al*, 1983; Triggiani *et al*, 1994; Bozza *et al*, 2011). It was only recently shown that ATGL provides precursors for lipid mediator production and modulates neutrophil immune responses *in vivo* (Dichlberger *et al*, 2014; Schlager *et al*, 2015; Riederer *et al*, 2017). However, it is not yet clear how LDs manage PUFA trafficking for lipid mediator production in mammalian cells and how they cooperate with the canonical PLA_2_-mediated pathways (Guijas *et al*, 2014; Jarc & Petan, 2020).

The trafficking of PUFAs between membranes and LDs is an emerging mechanism controlling PUFA oxygenation and lipid-induced oxidative damage, including ferroptotic cell death. On the one hand, the sequestration of PUFAs into LDs limits their availability for oxidation (Bailey *et al*, 2015; Jarc *et al*, 2018; Dierge *et al*, 2021). For instance, in fly embryos exposed to hypoxia, membrane lipid peroxidation and neuronal damage are prevented via a mechanism mediated by phospholipase D and diacylglycerol acyltransferase (DGAT) that diverts PUFAs from phospholipids into TAGs stored within LDs (Bailey *et al*, 2015). On the other hand, the proportion of (poly)unsaturated FAs released from LDs may determine membrane saturation, oxidative stress and cell fate (Ackerman *et al*, 2018; Jarc *et al*, 2018). Accordingly, in cancer cells exposed to exogenous PUFAs, LDs are enriched with PUFA-TAGs and their hydrolysis by ATGL promotes oxidative stress-dependent cell death (Jarc *et al*, 2018). Under these conditions, the transfer of unsaturated FAs from membrane phospholipids into LDs mediated by the human group X (hGX) sPLA_2_ balances phospholipid and TAG acyl-chain composition and prevents PUFA lipotoxicity (Pucer et al, 2013; Jarc *et al*, 2018). Based on these findings, we hypothesize that LD turnover modulates membrane PUFA content and affects PLA_2_-mediated lipid mediator production.

Here, we investigated the crosstalk between different molecular pathways providing PUFAs for lipid mediator production in human cancer cells. We show that the incorporation of PUFAs into TAGs and their subsequent release via lipolysis are essential for production of mitogenic lipid mediators. We demonstrate that LDs integrate several PUFA trafficking pathways and act as central hubs that control PLA_2_-driven lipid mediator production.

Furthermore, we provide evidence for a pathophysiological relevance of these findings by showing that inhibition of TAG synthesis impairs the production of mitogenic lipid mediators and reduces cancer cell proliferation in vitro and tumour growth in vivo.

## Results

### Membrane phospholipid hydrolysis by hGX sPLA_2_ leads to enrichment of LDs with long-chain PUFA-TAGs

We have shown previously that hGX sPLA_2_ releases various unsaturated FAs from adherent cells, including oleic acid (C18:1n–9; OA) and PUFAs (Jarc *et al*, 2018), and induces LD accumulation in several breast cancer cell lines (Pucer *et al*, 2013; Jarc *et al*, 2018). Recombinant hGX sPLA_2_ also induced LD accumulation in other cancer and immortalised non-tumorigenic cell lines (Fig. 1A, B), suggesting that its effects on LD metabolism are not limited to specific cell types. Here we asked whether hGX sPLA_2_ alters TAG acyl chain composition in LDs of MDA-MB-231 cells (Fig. 1C), a cell line showing enhanced proliferation and resistance to starvation upon treatment with hGX sPLA_2_ (Pucer *et al*, 2013). Cells pre-incubated with [^14^C]-OA readily incorporated the radiolabelled FA into phospholipids and TAGs, while hGX sPLA_2_ treatment specifically increased the abundance of TAGs containing [^14^C]-OA in various growth conditions (Fig. 1D; Supp. Fig. 1A, B). Lipidomic LC/MS analyses of TAG species extracted from hGX sPLA_2_-treated serum-fed (Fig. 1E–H, J, K; Supp. Fig. 1C, D) and serum-starved cells (Fig. 1F, I) revealed an unexpected enrichment of TAG species with long-chain (Fig. 1J) and highly unsaturated FAs (Fig. 1K). This suggests that among the unsaturated FAs released by hGX sPLA_2_, the polyunsaturated species are preferentially redistributed from membrane phospholipids into LDs. Experiments with serum-starved cells (Fig. 1D, F, I) demonstrated that serum lipoproteins, which are major targets for the enzyme (Guillaume *et al*, 2015; Jarc *et al*, 2018), are not required for hGX sPLA_2_-induced enrichment of TAGs with PUFAs. This observation suggests a direct action of hGX sPLA_2_ on the plasma membrane leading to enrichment of LDs with PUFA-containing TAGs.

**Figure 1.**
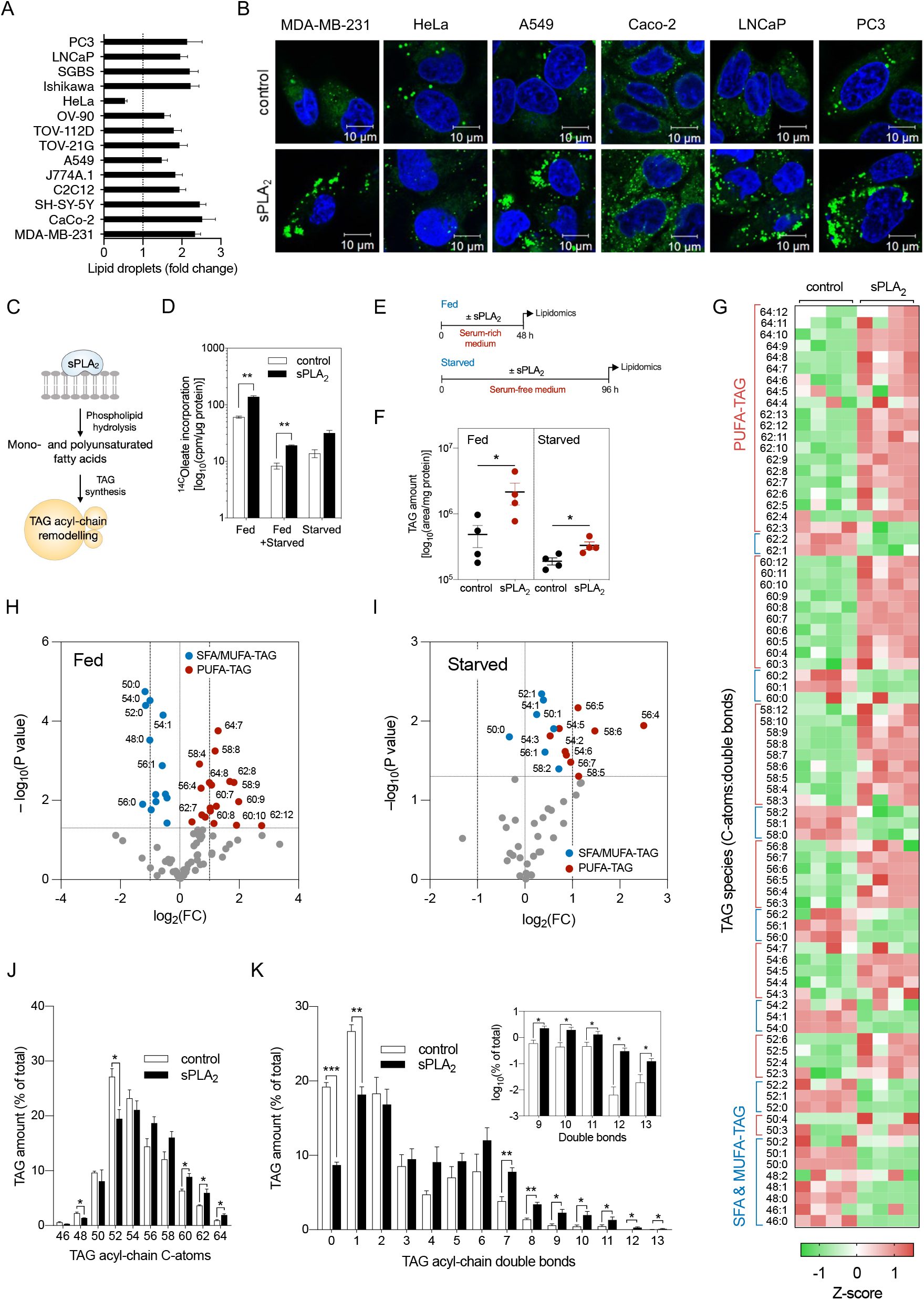
sPLA_2_ promotes enrichment of LDs with long-chain PUFA-containing triglycerides. (**A,B**)LD levels in control cells and cells treated with 10 nM recombinant hGX sPLA_2_ for 48 h in serum-rich medium. (**A**) Neutral lipid content was quantified by Nile Red staining and flow cytometry (n = 3 independent experiments; 20000 cells per treatment). (**B**) Representative live-cell confocal microscopy images of LDs stained with BODIPY 493/503 (green) and nuclei with Hoechst 33342 (blue). (**C**) Diagram illustrating the hypothesis that unsaturated fatty acids released through sPLA_2_ membrane hydrolysis are incorporated into triacylglycerols (TAGs) and lead to lasting changes in LD TAG acyl-chain composition. (**D**) hGX-sPLA_2_-induced changes in the incorporation of radiolabelled oleate into cellular TAGs in MDA-MB-231 cells grown in complete medium for 24 h (Fed), in complete medium for 24 h followed by 96 h of serum starvation (Fed + Starved), and in serum-free medium for 96 h (Starved) (n = 4 independent experiments). (**E**) Diagram illustrating the experimental treatments used for lipidomic analysis in (F)–(K). (**F–K**) TAG lipidomic analyses of MDA-MB-231 cells treated with recombinant hGX sPLA_2_ in complete medium for 48 h (Fed; F–H, J, K) or in the absence of serum for 96 h (Starved; F, I). Cell lysates were collected and analysed by UPLC/qTOF-MS (n = 4 independent experiments). (**G–I**) sPLA_2_-induced changes in the levels of individual TAG species presented as a representative z-score heat-map (G) and volcano plots (H, I) prepared by log_2_ data transformation and multiple t-test analyses (n = 4 independent experiments). Statistically significant changes (–log_10_(P value) > 1.30) in TAGs containing mostly saturated and mono-unsaturated FAs (SFA/MUFA-TAGs with 0–2 double bonds; blue) and those containing polyunsaturated FAs (PUFA-TAGs with 3–12 double bonds; red) are shown. (**J, K**) sPLA_2_-induced changes in relative levels of TAG species grouped by number of acyl-chain C-atoms (chain length) and double bonds (chain unsaturation). (A–K) Data are means ±SEM of at least three independent experiments. *, P <0.05; **, P <0.01; ***, P <0.001 (unpaired t-tests).

### ATGL-mediated LD breakdown is required for PGE_2_ production in serum-starved cells

Serum withdrawal induces LD breakdown in most cell types (Bosch *et al*, 2020). Given the hGX-sPLA_2_-induced enrichment of LDs with PUFAs, we next asked if starvation-induced breakdown of PUFA-rich LDs is sufficient to stimulate eicosanoid production. LD biogenesis was first stimulated with hGX sPLA_2_ (or with exogenous AA) in cells grown in serum-rich medium. Then the cells were serum-starved in the absence of these stimuli to induce LD breakdown (Fig. 2A; Supp. Fig. 2A). In comparison with untreated cells, the serum-starved cells pre-treated with hGX sPLA_2_ released more glycerol (Supp. Fig. 2C), an indicator of TAG lipolysis, and produced more prostaglandin (PG)E_2_ (Fig. 2B), a major AA-derived eicosanoid. Although HeLa cells were unique among cell lines in showing a net reduction in neutral lipid levels upon treatment with hGX sPLA_2_ (Fig. 1A, B; Supp. Fig. 2B), there was an increase in lipolysis (Supp. Fig. 2C) and PGE_2_ production (Fig. 2B) during serum starvation. This is indicative of an accelerated LD turnover in hGX-sPLA_2_-treated HeLa cells that leads to net reduction in LD levels and drives lipid mediator production. Similarly, pre-treatment with exogenous AA led to increased PGE_2_ production in starving MDA-MB-231 and HeLa cells (Supp. Fig. 2D). These observations indicate that LD breakdown drives PGE_2_ synthesis.

**Figure 2.**
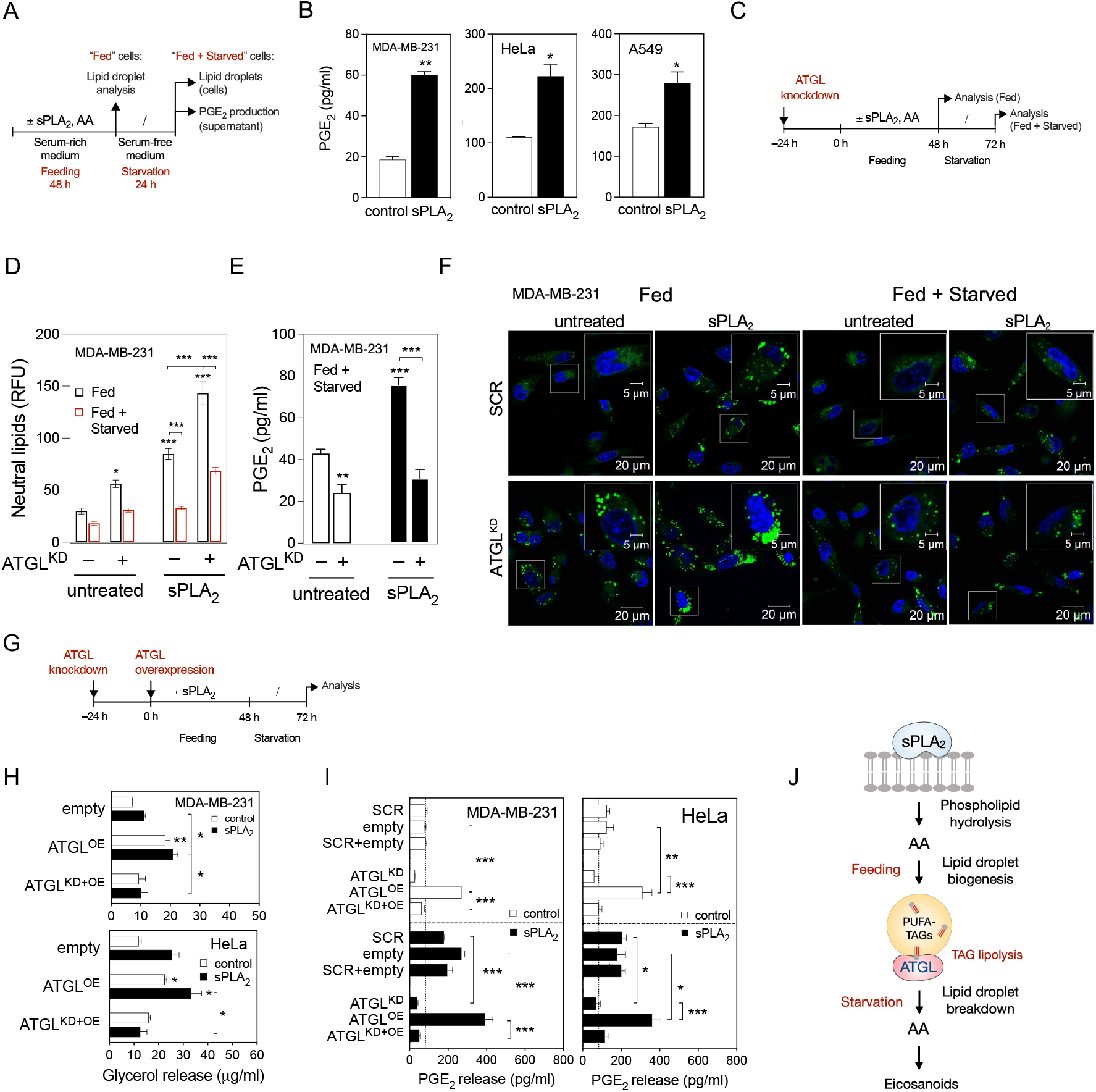
ATGL-mediated LD breakdown is required for PGE_2_ production in serum-starved cancer cells. (**A**) Diagram illustrating the experimental set-up used to load cells with LDs (Feeding) and then to induce their breakdown (Starvation). (**B**) PGE_2_ levels in cell supernatants of control and hGX-sPLA_2_-treated cells quantified by ELISA at the end of the starvation period. (**C**) Diagram illustrating the experimental conditions used in (D)–(F) and Supp. Fig. 1(E)–(O). (**D, E**) LD levels and PGE_2_ production in ATGL-silenced control and sPLA_2_-treated cells grown as shown in (C) and analysed at the beginning (Fed) and at the end of the serum starvation period (Fed + Starved). Neutral lipids were quantified by Nile Red staining and flow cytometry, PGE_2_ was quantified by ELISA. (**F**) Representative confocal microscopy images showing effects of ATGL depletion on cellular LD content in control and hGX-sPLA_2_-treated cells, under serum-rich (Fed) and serum-free (Fed + Starved) conditions. LDs and nuclei were stained using BODIPY 493/503 and Hoechst 33342, respectively. (**G**) Diagram illustrating the experimental set-up used in (H), (I) and Supp. Figure 1P, R. (**H, I**) Glycerol release and PGE_2_ production in ATGL-overexpressing serum-starved cells (both untreated and hGX sPLA_2_ pretreated), in comparison with cells co-transfected with ATGL-specific siRNAs (ATGL^KD+OE^), non-targeting siRNA (scrambled) and control plasmid (empty), grown as illustrated in (G). (**J**) Diagram illustrating the proposed model of LD-mediated eicosanoid production in cancer cells. Data are means ±SEM of two (H) or three (B–E, I) independent experiments. *, P <0.05; **, P <0.01; ***, P <0.001 (two-way ANOVA with Tukey (I) or Bonferroni (D, E, H) adjustment; unpaired t-tests (B)).

We next investigated if ATGL, the rate-limiting enzyme in intracellular TAG lipolysis, is responsible for LD breakdown and PGE_2_ biosynthesis observed during starvation. ATGL silencing using siRNA in MDA-MB-231 and HeLa cells (Fig. 2C; Supp. Fig. 2E) reduced glycerol release (Supp. Fig. 2F, G), increased LD abundance (Fig. 2D, F; Supp. Fig. 2H, J) and reduced basal and hGX-sPLA_2_-induced PGE_2_ production during serum starvation (Fig. 2E; Supp. Fig. 2I). ATGL deficiency also attenuated LD breakdown and suppressed the production of PGE_2_ in MDA-MB-231 cells pre-treated with exogenous AA (Supp. Fig. 2N, O). However, although ATGL knockdown increased LD abundance also in A549 cells (Supp. Fig. 2E; Supp. Fig. 2K, M), it unexpectedly augmented glycerol release (Supp. Fig. 2F, G) and failed to suppress basal or hGX-sPLA_2_-induced PGE_2_ production (Supp. Fig. 2L). In contrast to the effects of ATGL silencing, ATGL overexpression in the breast and cervical cancer cells (Fig. 2G; Supp. Fig. 2P) stimulated glycerol release (Fig. 2H) and enhanced PGE_2_ production (Fig. 2I), albeit without significantly altering neutral lipid levels (Supp. Fig. 2R). The effects of ATGL overexpression were fully reversed in the presence of ATGL-targeting siRNA (Fig. 2H, I). Collectively, these data demonstrate that LD breakdown via ATGL is required for basal, AA-induced and hGX-sPLA_2_-stimulated PGE_2_ production (Fig. 2J) in MDA-MB-231 and HeLa cells, but not in the A549 cells.

### ATGL-mediated release of ω-3 and ω-6 PUFAs from LDs drives lipid mediator production

The data presented above demonstrate that transient storage of AA within LD TAGs followed by AA release from TAGs by ATGL are intermediate steps in basal and hGX-sPLA_2_-stimulated production of PGE_2_. To find out if ATGL-mediated TAG lipolysis promotes the production of a wider range of eicosanoids and related lipid mediators (which might also derive from other PUFAs), we performed targeted liquid chromatography-tandem mass spectrometry-based metabololipidomics to examine the effects of ATGL silencing on the lipid mediator profiles of untreated and hGX-sPLA_2_-treated cells. Here, pre-treatment of breast cancer cells with hGX sPLA_2_ stimulated the starvation-induced production of numerous lipid mediators, biosynthesised from different PUFAs by various enzymatic pathways (Fig. 3A–C, Supp. Fig. 3). hGX sPLA_2_ promoted the synthesis of eicosapentaenoic acid (EPA)-derived lipid mediators, as well as the production of AA-derived and DHA-derived products. ATGL depletion suppressed the release of all of the hGX-sPLA_2_-induced lipid mediators (Fig. 3A–C, Supp. Fig. 3). Furthermore, ATGL silencing strongly inhibited the basal, hGX-sPLA_2_-independent production of lipid mediators, including PGE_2_. Starving cells pre-treated with hGX sPLA_2_ also released more unesterified PUFAs, including AA, EPA and DHA, which was prevented by ATGL silencing (Fig. 3D). This suggested that a portion of hGX sPLA_2_-released PUFAs is not used for lipid mediator production or incorporated into membranes, but is rather released from cells through lipolysis by ATGL. Overall, these observations suggest that ATGL controls the availability of PUFAs for lipid mediator synthesis (Fig. 3E).

**Figure 3.**
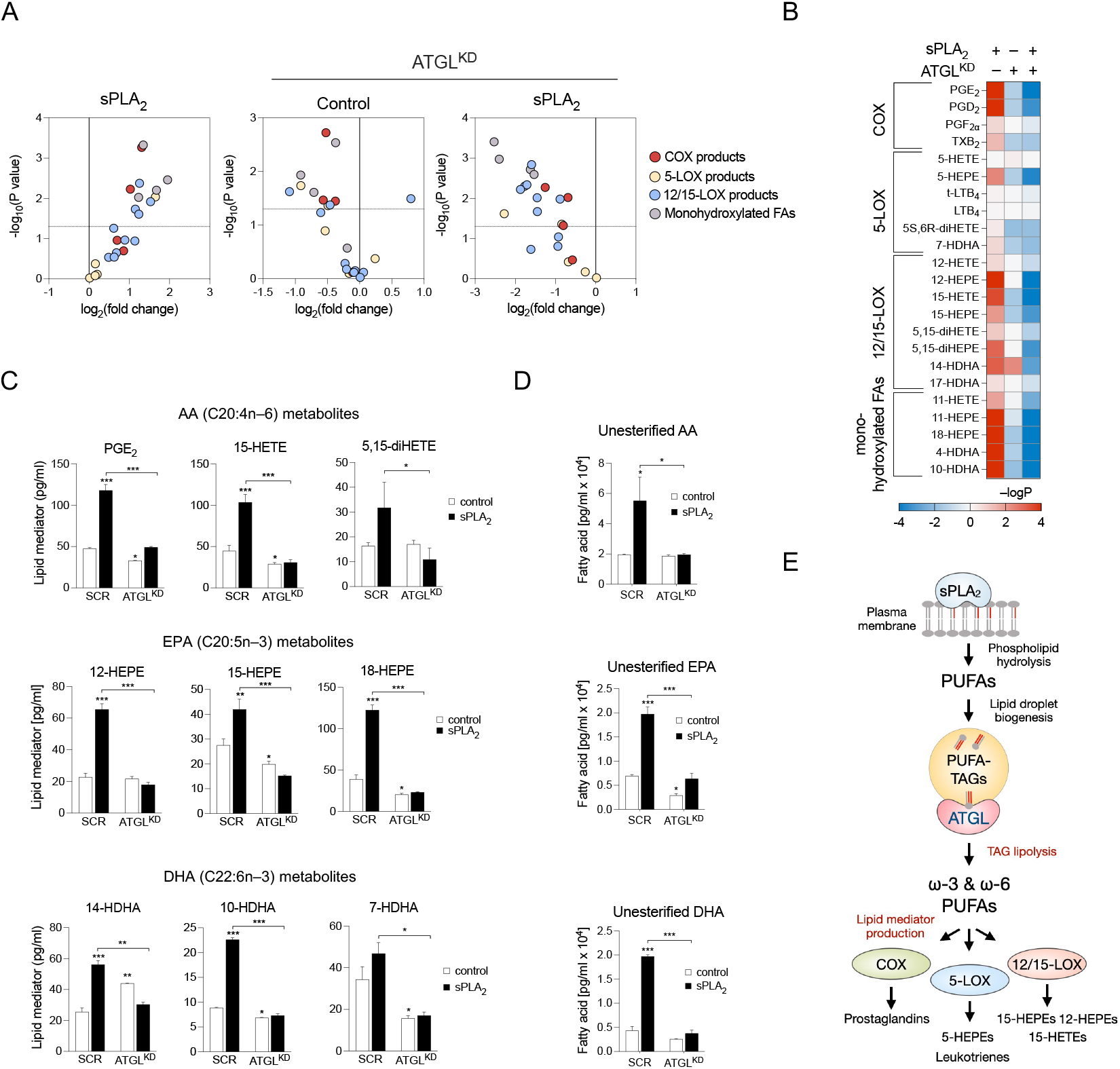
ATGL-mediated lipolysis drives production of a wide spectrum of bioactive lipid mediators. (**A–D**) UPLC-MS-MS analysis of lipid mediators released from serum-starved MDA-MB-231 cells pre-treated with hGX sPLA_2_, and from ATGL-depleted cells without and with hGX sPLA_2_ pre-treatment, presented as volcano plots (A), a heat map (B), and individual graphs (C, D). Cells were treated as shown in Fig. 2C. Volcano plots were prepared using log_2_-transformed fold-change values and multiple t-test analysis, and the heat map by −logP data transformation and two-way ANOVA with Sidak adjustment (n = 3 independent experiments). (**E**) Diagram illustrating the involvement of hGX sPLA_2_ and ATGL in the production of a wide range of PUFA-derived cyclooxygenase (COX) and lipoxygenase (LOX) signalling molecules. Data are means ±SEM of three independent experiments. *, P <0.05; **, P <0.01; ***, P <0.001 (two-way ANOVA with Sidak adjustment).

### Inhibition of DGAT-mediated LD biogenesis impairs PGE_2_ production

To examine whether the incorporation of PUFAs into TAGs is a prerequisite for the conversion of PUFAs into lipid mediators, we inhibited TAG synthesis using a combination of specific inhibitors of DGAT1 and DGAT2 (DGATi), which mediate the final committed step in TAG synthesis, and followed changes in LD turnover and lipid mediator production. Combined inhibition of both DGATs was necessary because individual inhibition of DGAT1 or DGAT2 did not reduce neutral lipid levels in hGX-sPLA_2_-treated cells (Supp. Fig. 4A). Combined DGAT1 and DGAT2 inhibition during serum feeding strongly suppressed basal, hGX-sPLA_2_-induced and exogenous AA-induced LD accumulation (Fig. 4A, B, D). There was an almost complete depletion of LDs in serum-fed DGATi-treated cells (Fig. 4B), which allowed only a minimal breakdown of residual LDs during the subsequent serum starvation (Fig. 4A). Notably, cells depleted of LDs displayed low basal PGE_2_ release and did not increase PGE_2_ production upon stimulation with either hGX sPLA_2_ or exogenous AA (Fig. 4C, E). These results were corroborated by lipidomic analyses, which showed that combined inhibition of DGAT1 and DGAT2 suppresses hGX sPLA_2_-induced production of several AA- and EPA-derived lipid mediators (Fig. 4F; Supp. Fig. 4B), reduces exogenous AA-induced production of prostaglandins (Supp. Fig. 4C) and blunts the release of unesterified PUFAs from starving MDA-MB-231 cells (Fig. 4G). Notably, DGATi treatment abolished PGE_2_ production also in A549 cells (Fig. 4C), whereas ATGL depletion failed to do so (Supp. Fig. 2L), suggesting that TAG synthesis precedes eicosanoid production in these cells as well. Therefore, DGAT-mediated TAG biosynthesis under serum-rich conditions is a prerequisite for lipid mediator production in serum-starved cells.

**Figure 4.**
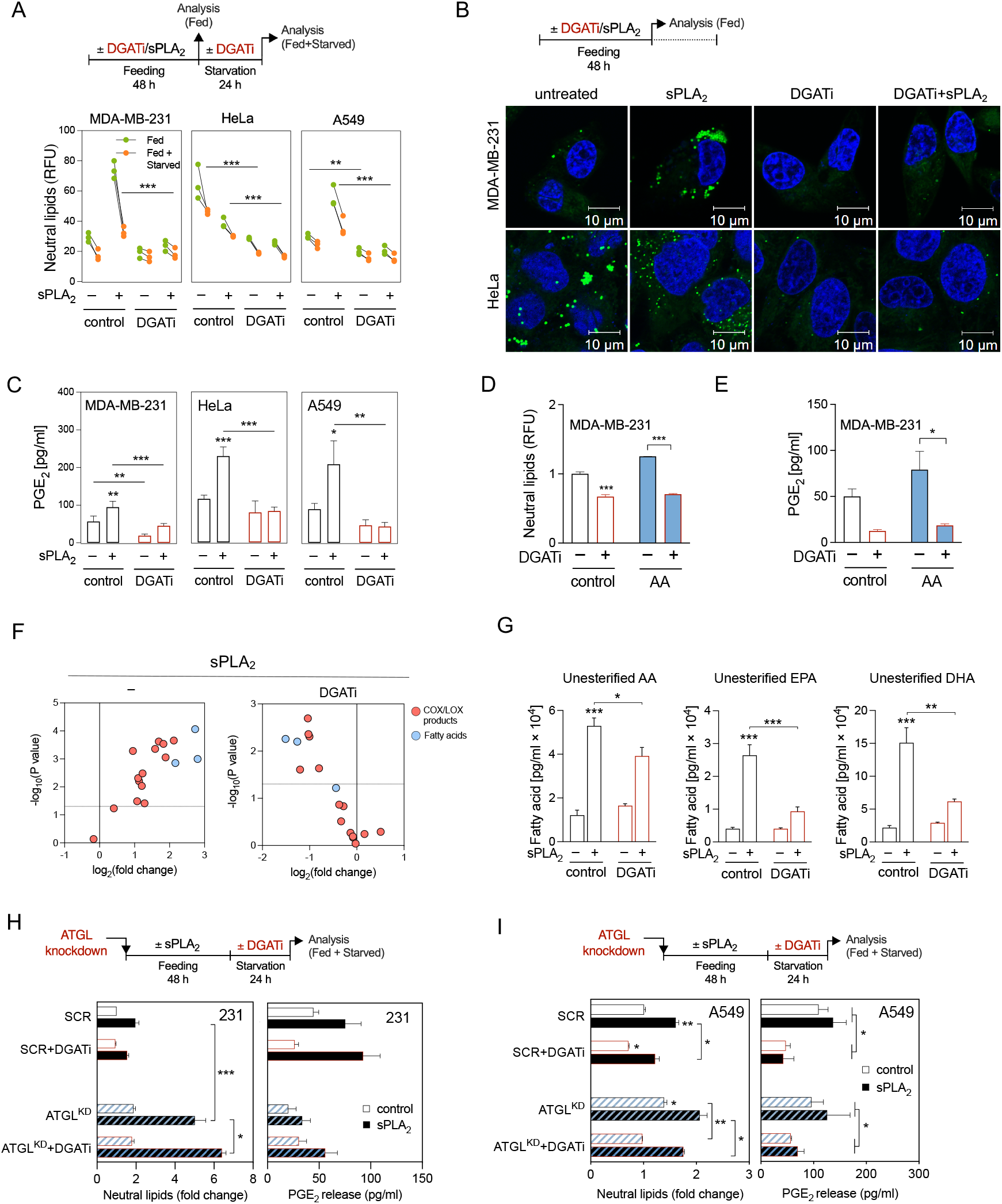
DGAT-mediated LD biogenesis is required for lipid mediator production in serum-starved cancer cells. (**A**) Diagram illustrating the experimental conditions used (top) and cellular neutral lipid content before (Fed) and after (Fed + Starved) serum starvation in cells treated with DGAT1 (T863) and DGAT2 (PF-06427878) inhibitors (DGATi), without and with stimulation of LD biogenesis by hGX sPLA_2_ during serum sufficiency. (**B**) Representative confocal microscopy images of live cells under nutrient-replete conditions and treated with DGAT inhibitors in the absence and presence of hGX sPLA_2_. LDs and nuclei were visualised using BODIPY 493/503 and Hoechst 33342 staining, respectively, and confocal microscopy. (**C**) DGATi-induced changes in PGE_2_ production in serum-starved cancer cells (Fed + Starved), without and with additional stimulation of LD biogenesis by hGX sPLA_2_ pre-treatment. Cells were treated according to (A). (**D, E**) DGATi-induced changes in neutral lipids (D) and PGE_2_ production (E) in serum-starved MDA-MB-231 cancer cells (Fed + Starved), without and with additional stimulation of LD biogenesis by arachidonic acid (AA) pre-treatment. Cells were treated according to (A). (**F, G**) DGATi-induced changes in the profiles of lipid mediators and fatty acids released from serum-starved MDA-MB-231 cells treated with hGX sPLA_2_, without and with treatment with an equimolar mix of DGAT1 and DGAT2 inhibitors according to (A). Volcano plots (F) show significant changes (−log_10_(P value) >1.30) in individual lipids between hGX sPLA_2_-treated *versus* control cells (left) and hGX sPLA_2_-treated *versus* hGX sPLA_2_- and DGATi-treated (right) cells and were prepared using log_2_-transformed fold-change values and multiple t-test analysis (n = 4 independent experiments). (**H, I**) Diagrams illustrating the experimental conditions used (top) and changes in PGE_2_ production induced by DGATi treatments during serum starvation in control (SCR) and ATGL-depleted (ATGL^KD^) MDA-MB-231 (H) and A549 (I) cells, without and with hGX sPLA_2_ pre-treatment. (A, D, H, I) Neutral lipid content was quantified by Nile Red staining and flow cytometry. (C, E, H, I) PGE_2_ levels were determined in cell supernatants as described in Methods. (F, G) UPLC-MS/MS analysis of lipid mediators and FAs in cell supernatants was performed as described in Methods. Data are means ±SEM of at least three (A, C, D, E, H, I) or four (F, G) independent experiments. *, P <0.05; **, P <0.01; ***, P <0.001 (two-way ANOVA with Tukey (A, H, G, I) or Bonferroni (C, D, E) adjustment).

Besides promoting LD breakdown, serum removal in MDA-MB-231 cells stimulates the transcription of lipogenic genes, including the sterol-regulatory element binding 1 protein (SREBP-1) transcription factor and key enzymes involved in FA synthesis (Pucer *et al*, 2013). In accordance with ongoing lipogenesis, treatment of serum-starved cells with hGX sPLA_2_ resulted in a net increase in TAG levels (Fig. 1F). To examine whether TAG synthesis occurring during serum starvation regulates eicosanoid production, control and ATGL-depleted cells were treated with DGAT1 and DGAT2 inhibitors during serum feeding or during serum starvation. As expected, DGAT inhibition during serum feeding fully depleted MDA-MB-231 and HeLa cells of LDs, thereby abolishing the effects of hGX sPLA_2_ and ATGL silencing on LD abundance (Supp. Fig. 4D). In contrast, DGAT inhibition during serum starvation did not affect LD abundance in these two cell lines (Fig. 4H; Supp. Fig. 4E), nor did it affect PGE_2_ release in MDA-MB-231 cells (Fig. 4H). However, DGAT inhibition in serum-starved A549 cells reduced LD abundance and suppressed PGE_2_ production (Fig. 4I). This was observed in both control and in ATGL-deficient A549 cells indicating that the build-up of TAG stores during starvation drives eicosanoid biosynthesis via an ATGL-independent mechanism. Accordingly, DGAT inhibition during both serum feeding and starvation was necessary for full suppression of PGE_2_ production in A549 cells (Supp. Fig. 4F).

Collectively, these results demonstrate that DGAT-mediated TAG synthesis and LD turnover are required for lipid mediator production. ATGL-mediated breakdown of pre-existing DGAT-induced LDs is the predominant mechanism of LD-driven eicosanoid production in the serum-starved MDA-MB-231 breast and HeLa cervical cancer cells. On the contrary, A549 lung cancer cells employ TAG synthesis during starvation and ATGL-independent mechanisms to support eicosanoid production.

### cPLA_2_α cooperates with ATGL and depends on LD turnover to drive eicosanoid production

Given the well-accepted role of cPLA_2_α in providing AA for eicosanoid biosynthesis, we next asked whether cPLA_2_α participates in LD-driven lipid mediator production. We speculated that at least three main scenarios are possible (Fig. 5A): (a) cPLA_2_α-induced incorporation of phospholipid-derived AA into TAGs, followed by AA release from LDs by ATGL; (b) ATGL-dependent transfer of TAG-derived AA into phospholipid pools, which are then accessed by cPLA_2_α; and (c) a lipid-droplet-independent action of cPLA_2_α on membrane phospholipids.

**Figure 5.**
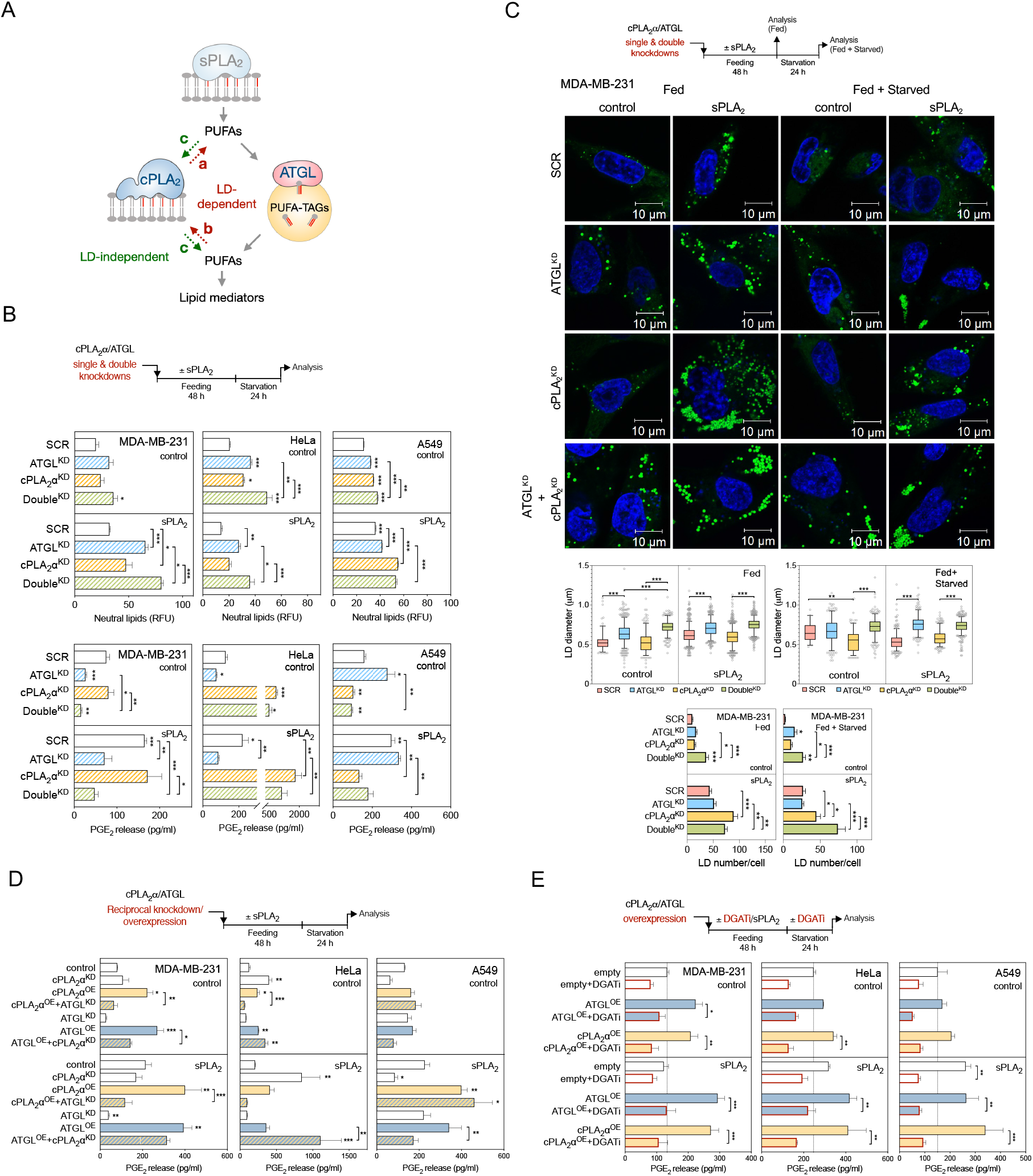
cPLA_2_α depends on LD turnover to drive lipid mediator production. (**A**) Scheme illustrating hypothetical models of interplay between cPLA_2_α and LDs in providing PUFAs for lipid mediator production. (**B**) Changes in neutral lipid content and PGE_2_ production induced by ATGL (ATGL^KD^) and cPLA_2_α (cPLA_2_α^KD^) single and double (Double^KD^) knockdowns, in comparison with control siRNA-treated cells (SCR), without and with stimulation of LD biogenesis by hGX sPLA_2_ pre-treatment. (**C**) Diagram illustrating the experimental conditions used (top), representative live-cell confocal microscopy images (middle) and image analysis (bottom) of LDs in ATGL (ATGL^KD^) and cPLA_2_α (cPLA_2_α^KD^) single and double (Double^KD^) knockdown MDA-MB-231 cells, in comparison with non-targeting control siRNA-treated cells (SCR), without and with stimulation of LD biogenesis by hGX sPLA_2_ pre-treatment. LDs were stained with BODIPY 493/503 (green) and nuclei with Hoechst 33342 (blue) and images analysed using ImageJ and the LD Counter Plugin. Box plots are showing changes in LD diameters and bar plots changes in LD numbers per cell in serum-fed (Fed) and serum-starved (Starved) cells. Data are geometric means (diameter analysis) or means (number analysis) ±SEM (n >40 cells/sample) of two independent experiments. (**D**) PGE_2_ production in cells with reciprocal knockdown/overexpression of cPLA_2_α and ATGL. Cells were reverse transfected with siRNAs specific for ATGL (ATGL^KD^) and/or cPLA_2_α (cPLA_2_α^KD^), then forward transfected with ATGL-encoding (ATGL^OE^) and/or cPLA_2_α-encoding (cPLA_2_α^OE^) plasmids, without and with pre-treated with hGX sPLA_2_, as illustrated in the scheme (top). In controls (control), non-targeting siRNA reverse transfections were combined with backbone (‘empty’) vector forward transfections. (**E**) DGAT inhibition (DGATi)-induced changes in PGE_2_ production in serum-starved control cells (empty) and in cells overexpressing ATGL (ATGL^OE^) or cPLA_2_α (cPLA_2_α^OE^), without and with additional stimulation of LD biogenesis by hGX sPLA_2_ pre-treatment. Neutral lipid content was quantified by Nile Red staining and flow cytometry (B), PGE_2_ levels were determined in cell supernatants using ELISA (B, D, E). Data are means ±SEM of two (B, A549 cells) or at least three independent experiments *, P <0.05; **, P <0.01; ***, P <0.001 (nested one-way ANOVA with Sidak adjustment (C, box plots), two-way ANOVA with Tukey (B, C, E) or Dunnet (D) adjustment).

To examine these possibilities, we first used siRNA to deplete cells of cPLA_2_α alone or in combination with ATGL (Fig. 5B; Supp. Fig. 5A, B). cPLA_2_α deficient cancer cells had slightly increased neutral lipid levels (Fig. 5B; Supp. Fig. 5C), but contained a significantly higher number of LDs (Fig. 5C; Supp. Fig. 5E, F). This finding was particularly evident in ATGL-deficient and hGX sPLA_2_ pre-treated starving cells and suggests a modulatory role of cPLA_2_α in LD turnover. In line with its major role in TAG lipolysis, the predominant effect of ATGL silencing on LD morphology was an increase in LD diameters, along with some minor changes in the number of LDs per cell (Fig. 5C; Supp. Fig. 5E, F). Most notably, cPLA_2_α depletion in A549 cells, but not ATGL silencing, abolished basal and hGX-sPLA_2_-induced eicosanoid release (Fig. 5B). On the contrary, while ATGL knockdown lowered PGE_2_ production in MDA-MB-231 cells, cPLA_2_α silencing did not alter PGE_2_ levels. Of note, the silencing of cPLA_2_α in HeLa cells led to a marked increase in PGE_2_ release, which was partially reduced upon depletion of both ATGL and cPLA_2_α (Fig. 5B). These data suggested that cPLA_2_α modulates LD turnover and that its contribution to PGE_2_ production during serum starvation is cell type-specific.

To find out more about the interplay between cPLA_2_α and ATGL, we asked how reciprocal overexpression of one and silencing of the other might affect LD turnover and PGE_2_ production (Fig. 5D). In all three cell lines, the overexpression of ATGL or cPLA_2_α had a minimal effect on neutral lipid levels (Supp. Fig. 5D, G, H), but both significantly augmented PGE_2_ production (Fig. 5D). ATGL silencing blocked PGE_2_ production induced by overexpression of cPLA_2_α in MDA-MB-231 and HeLa cells, whereas cPLA_2_α silencing only partially reduced ATGL-overexpression-driven eicosanoid production in MDA-MB-231 cells, and even increased it in HeLa cells. On the contrary, in A549 cells, ATGL knockdown did not affect cPLA_2_α-induced PGE_2_ production, but silencing of cPLA_2_α reduced PGE_2_ release from ATGL-overexpressing cells (Fig. 5D). Taken together, these data suggested that ATGL is the main enzyme in the provision of PUFAs for eicosanoid production in MDA-MB-231 and HeLa cells, whereby cPLA_2_α stimulation of eicosanoid production depends on ATGL. In contrast, in A549 cells, cPLA_2_α has a dominant role in PGE_2_ production, which is independent of ATGL, but is still associated with changes in LD turnover.

To examine if TAG synthesis is involved in cPLA_2_α-driven lipid mediator production, cPLA_2_α and ATGL were overexpressed in cells depleted of LDs using DGAT inhibitors (Fig. 5E; Supp. Fig. 5H). As expected, TAG synthesis was necessary for ATGL stimulation of eicosanoid production (Fig. 5E). However, in all three cell lines, cPLA_2_α-overexpression-induced eicosanoid production was also completely abolished upon DGAT inhibition (Fig. 5E). Therefore, LDs are required for cPLA_2_α-driven lipid mediator production. cPLA_2_α and ATGL appear to have cell-type-specific roles that are either cooperative or complementary for both LD turnover and eicosanoid production. In the breast and cervical cancer cells, deficiency of ATGL impairs cPLA_2_α-induced PGE_2_ production, which suggests the possibility that the transfer of AA from TAGs to phospholipids is a prerequisite for the action of cPLA_2_α.

### ATGL and cPLA_2_α have complementary roles in TAG and phospholipid acyl-chain remodelling

To examine how hGX sPLA_2_, ATGL and cPLA_2_α affect PUFA trafficking between the phospholipid and TAG pools in MDA-MB-231 cells during serum starvation, we performed single and double knockdowns of ATGL and cPLA_2_α followed by lipidomic analyses comparing the lipid compositions of cells first pre-treated with hGX sPLA_2_ during nutrient sufficiency and subsequently serum-starved for 0 h, 3 h or 24 h (Fig. 6A).

**Figure 6.**
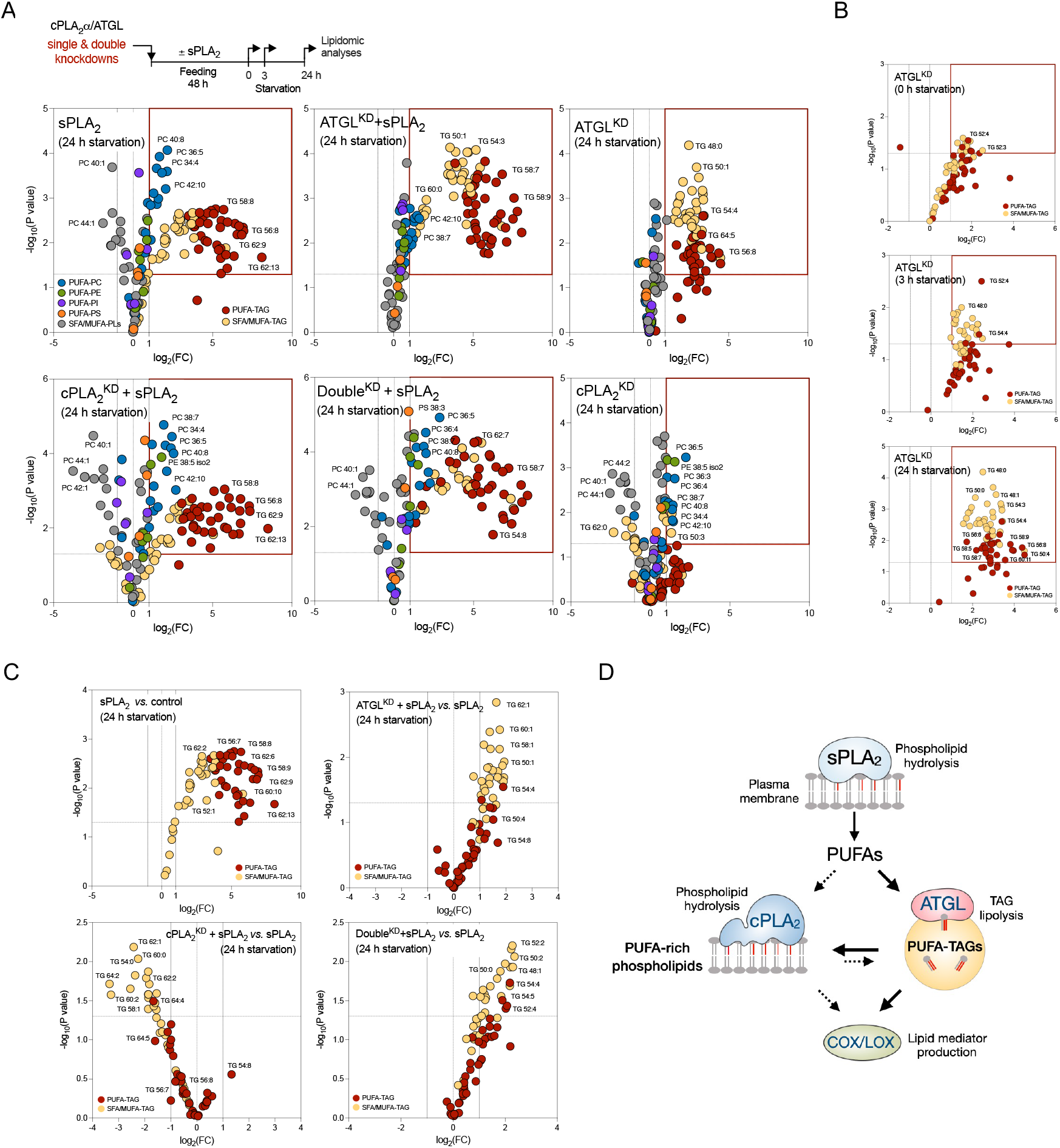
ATGL and cPLA_2_α cooperatively modulate PUFA trafficking between triglycerides and membrane phospholipids. (**A–C**) Untargeted lipidomic analysis of phospholipids and triglycerides (TAGs) in serum-starved MDA-MB-231 cells depleted of ATGL (ATGL^KD^), cPLA_2_α (cPLA_2_α^KD^), or both (Double^KD^), without and with hGX sPLA_2_ pre-treatment under serum-rich conditions, and grown as shown in the diagram (A). Volcano plots show significant changes (−log_10_(P value) >1.30) in individual lipids between each treatment condition *versus* control cells (unless otherwise indicated), and were prepared by log_2_ fold-change (FC) data transformation and multiple t-test analysis (n=3 independent experiments). TAGs and phospholipids (PLs) containing saturated and mono-unsaturated acyl chains (SFA/MUFA-TAGs, SFA/MUFA-PLs, with 0–3 and 0–2 double bonds, respectively) and those containing polyunsaturated FAs (PUFA-TAGs, PUFA-PLs, with at least 4 and 3 double bonds, respectively) are colour-coded as indicated. TG, triglyceride; PC, phosphatidylcholine; PE, phosphatidylethanolamine; PI, phosphatidylinositol, PS, phosphatidylserine. (**D**) Schematic illustration of the predominant pathways involved in LD-mediated PUFA trafficking between the membrane phospholipid and TAG pools in serum-starved cancer cells.

First, hGX sPLA_2_ treatment led to enrichment of both TAGs and phospholipids with PUFAs (Fig. 6A; Supp. Fig. 6A; Supp. Fig. 7A). hGX-sPLA_2_-induced PUFA-TAG enrichment was seen for serum-fed MDA-MB-231 cells, and it persisted upon hGX sPLA_2_ and serum withdrawal. Importantly, the levels of PUFA-containing phospholipids (PUFA-PLs) progressively increased during the course of serum starvation (Supp. Fig. 6A). These data suggest that a significant portion of sPLA_2_-released PUFAs that are incorporated into TAGs in serum-fed cells are gradually released via lipolysis during serum starvation and are re-esterified into phospholipids.

Second, ATGL silencing led to an increased abundance of most TAG species, but also altered phospholipid composition (Fig. 6A; Supp. Fig. 6A). The retention of various TAG species within LDs due to ATGL deficiency is in agreement with the general lack of TAG acyl-chain specificity of ATGL (Eichmann et al, 2012). During serum starvation, ATGL depletion caused a gradual increase of numerous PUFA-TAGs (Fig. 6B), which confirmed that ATGL hydrolyses PUFA-TAGs in starving cells. Importantly, ATGL deficiency reverted the elevation of PUFA-PLs induced by hGX sPLA_2_ pre-treatment (Supp. Fig. 6A). This is consistent with a reduced flux of PUFAs from TAGs into phospholipids due to ATGL depletion. Thus, ATGL hydrolyses various TAG species, including numerous PUFA-TAGs, and provides PUFAs for esterification into phospholipids.

Third, depletion of cPLA_2_α predominantly affected phospholipid (Fig. 6A; Supp. Fig. 6A), but also altered TAG composition (Fig. 6C). Numerous PUFA-PC and PUFA-phosphatidylethanolamine (PUFA-PE) species were progressively elevated in cPLA_2_α-deficient cells over the course of the serum starvation (Supp. Fig. 6A), which suggested that these lipids are targeted by cPLA_2_α. The decreased abundance of PUFA-PLs observed in ATGL-deficient cells was reversed in double-knockdown cells (Fig. 6A; Supp. Fig. 6B), which suggested that ATGL-derived PUFAs are incorporated into a phospholipid pool that is targeted by cPLA_2_α. A LION/web lipid ontology analysis suggested that the increased content of PUFA-PLs in cPLA_2_α-deficient cells leads to significant changes in membrane biophysical properties, including increased lateral diffusion, reduced bilayer thickness and a lower transition temperature (Supp. Fig. 7B, C). Furthermore, cPLA_2_α silencing did not affect hGX-sPLA_2_-induced PUFA-TAG enrichment (Fig. 6A), but it reduced the content of SFA/MUFA-TAGs (Fig. 6C). This effect was fully reversed in double-knockdown cells, suggesting that cPLA_2_α depletion alters TAG composition in an ATGL-dependent manner. Therefore, ATGL promotes the transfer of PUFAs from TAGs into a phospholipid pool that is targeted by cPLA_2_α, which in turn reciprocally modulates the TAG pool.

Collectively, our results suggest that hGX-sPLA_2_-liberated PUFAs are first and predominantly incorporated into TAGs of growing LDs, and are then redistributed into phospholipids upon TAG lipolysis by ATGL, particularly during serum starvation (Fig. 6D). Some of the ATGL-released PUFAs are directly used for eicosanoid production (i.e., independent of cPLA_2_α), while some are re-esterified into phospholipids and can be then mobilised by cPLA_2_α for the production of eicosanoids.

### LDs drive cancer cell proliferation by promoting eicosanoid production

Given that PGE_2_ and other eicosanoids are mitogenic factors that promote tumour growth (Wang & DuBois, 2010), we next asked whether LD turnover by DGAT and ATGL affects cancer cell proliferation and possibly mediates the proliferative effect of hGX sPLA_2_ (Surrel *et al*, 2009; Pucer *et al*, 2013). The induction of MDA-MB-231 cell proliferation by hGX sPLA_2_ was fully blocked by DGAT1 inhibition and by concurrent inhibition of both DGAT1 and DGAT2; however, DGAT2 inhibition alone had no effects (Fig. 7A). Inhibition of DGAT1, but not DGAT2, also reduced the basal, hGX-sPLA_2_-independent rate of breast cancer cell proliferation (Fig. 7A) and colony formation (Supp. Fig. 8A, B). As ATGL promotes lipid mediator production, we hypothesized that ATGL overexpression should induce cell proliferation in a COX/LOX-dependent manner. Indeed, alone or in combination with sPLA_2_ treatment, ATGL overexpression stimulated cancer cell proliferation (Fig. 7B, C), which was suppressed by the non-selective COX and LOX inhibitors indomethacin and nordihydroguaiaretic acid, respectively (Fig. 7B, C). Exogenous PGE_2_ stimulated MDA-MB-231 cell proliferation, confirming its mitogenic potency under the present conditions (Supp. Fig. 8C). The effect of PGE_2_ was suppressed by inhibition of DGAT1, but not by inhibition of DGAT2, which suggested a positive-feedback loop between eicosanoids and DGAT1-mediated TAG turnover. To examine the relevance of DGAT1-driven lipid mediator production for cancer cell proliferation *in vivo*, we treated mice bearing aggressive MDA-MB-231 xenografts with DGAT1 inhibitors, followed tumour growth and determined lipid mediator content in tumours. DGAT1 inhibitor treatments reduced tumour volume and prolonged the survival of mice (Fig. 7D, E). Furthermore, tumour samples of DGAT1 inhibitor-treated mice contained reduced levels of lipid mediators (Fig. 7F). Thus, DGAT1 inhibition supressed lipid mediator production and tumour growth leading to prolonged survival of mice bearing breast cancer xenografts. Altogether, these results with MDA-MB-231 cancer cells demonstrate that DGAT1-mediated PUFA incorporation into LDs and PUFA release via ATGL drive basal and hGX-sPLA_2_-induced production of COX- and LOX-derived lipid mediators that in turn stimulate cancer cell proliferation and tumour growth.

**Figure 7.**
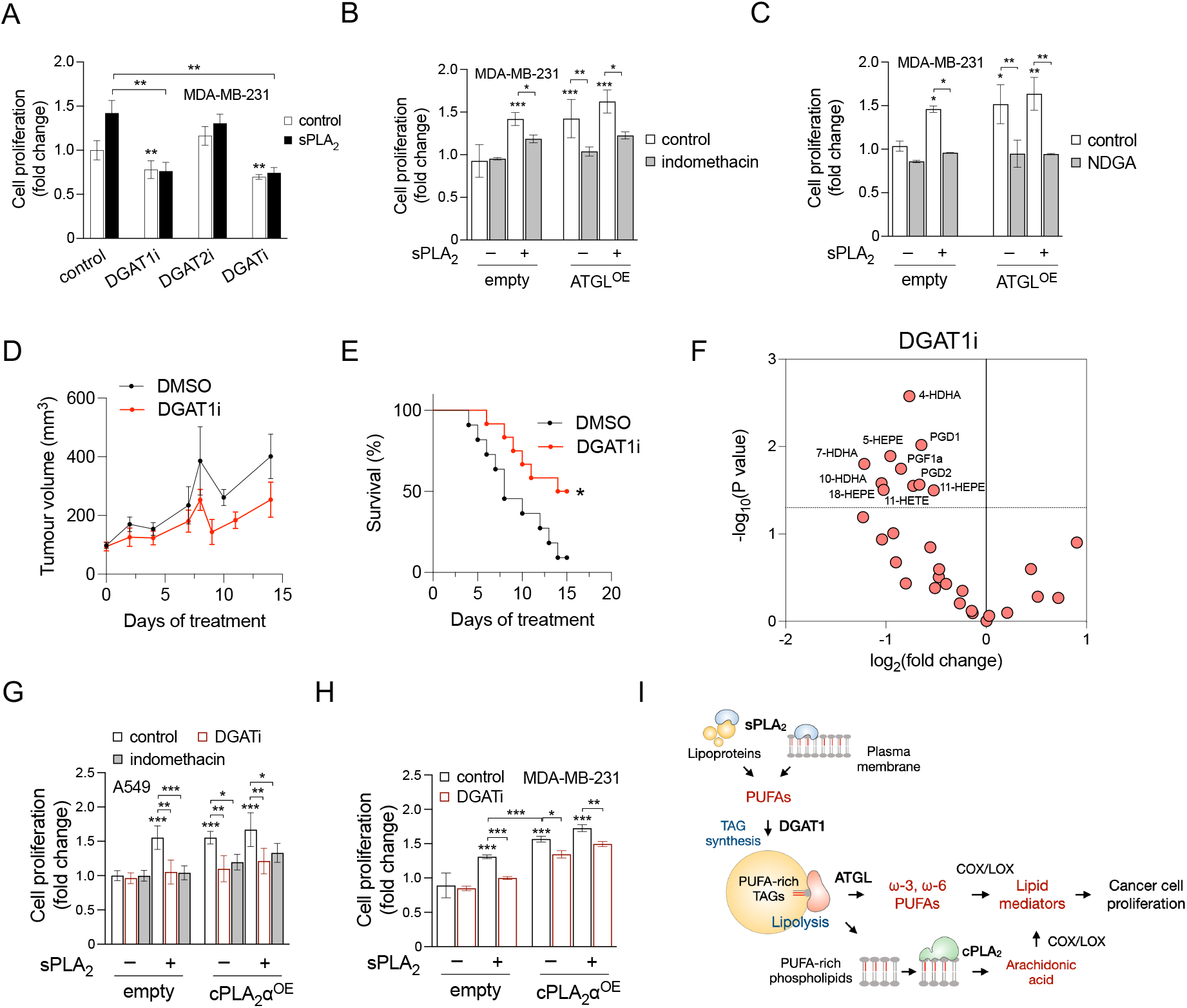
LD-mediated lipid mediator production promotes cancer-cell proliferation and tumor growth. (**A**) Proliferation of MDA-MB-231 cells treated with T863 (DGAT1i) and PF-06427878 (DGAT2i), or an equimolar mix of both DGAT inhibitors (DGATi), grown under nutrient-rich conditions in the absence and presence of hGX sPLA_2_. (**B, C**) Proliferation of MDA-MB-231 cells overexpressing ATGL (ATGL^OE^), treated with indomethacin (B) or nordihydroguaiaretic acid (NDGA) (C), in the absence and presence of hGX sPLA_2_. (**D**, **E**) Tumour growth (D) and corresponding Kaplan-Meier survival curves (E) of mice bearing MDA-MB-231 xenografts and treated daily with the DGAT1 inhibitor T863 or with 0.2% DMSO vehicle. Significance was determined by log-rank (Mantel-Cox) test (**F**) DGAT1-inhibitor-induced changes in the abundance of lipid mediators and PUFAs in tumour samples isolated from mice bearing MDA-MB-231 xenografts and treated daily with DGAT1 inhibitor T863 or with 0.2% DMSO vehicle. The volcano plot depicts significant changes (−log_10_(P value) >1.30) in individual lipids detected by UPLC-MS/MS in DGAT1i-treated *versus* vehicle-treated mice and was prepared using log_2_-transformed fold-change values and multiple t-test analysis (n = 4 independent experiments). (**G**) Proliferation of A549 cells overexpressing cPLA_2_α (cPLA_2_α^OE^) and grown in the absence and presence of DGATi, indomethacin and recombinant hGX sPLA_2_. (**H**) Proliferation of MDA-MB-231 cells overexpressing cPLA_2_α (cPLA_2_α^OE^) and grown in the absence and presence of DGATi and recombinant hGX sPLA_2_. Data are means ±SEM of three independent experiments. *, P <0.05; **, P <0.01; ***, P <0.001 (two-way ANOVA with Tukey (A), Bonferroni (B, C) or Sidak (G, H) adjustments).

cPLA_2_α-induced eicosanoid production promotes cancer cell proliferation and tumour growth (Leslie, 2015; Koundouros *et al*, 2020), although this activity has not yet been associated with LDs. As cPLA_2_α has a major role in LD-dependent eicosanoid production in A549 cells (Fig. 5B), we asked whether cPLA_2_α stimulates A549 cell proliferation in a DGAT- and COX-dependent manner. Indeed, cPLA_2_α-overexpression led to increased A549 cell proliferation, which was blocked by either DGAT or COX inhibitors (Fig. 8G), demonstrating that DGAT-mediated TAG synthesis and COX-mediated conversion of cPLA_2_α-released AA into lipid mediators are required for cPLA_2_α-stimulated A549 lung cancer cell proliferation. Exogenous hGX sPLA_2_ also stimulated A549 cell proliferation, which was suppressed by inhibition of DGAT and COX (Fig. 8G). Finally, in MDA-MB-231 cells, where DGAT-mediated TAG synthesis was required for cPLA_2_α-stimulated PGE_2_ production (Fig. 5E), cPLA_2_α overexpression resulted in higher proliferation rates, which were reduced by DGAT inhibition (Fig. 8H). Together, these data demonstrate that LD biogenesis controls both hGX-sPLA_2_- and cPLA_2_α-mediated cancer cell proliferation, which is driven by lipid mediators produced by COX and LOX enzymes.

## Discussion

In this study, we provide evidence that TAG turnover controls the production of a wide range of ω-3 and ω-6 PUFA-derived oxygenated lipid mediators. We show that the esterification of PUFAs into TAGs and their lipolytic release from LDs are necessary for PUFA entry into lipid mediator biosynthetic pathways. We demonstrate that ATGL liberates ω-3 and ω-6 PUFAs from TAGs, alters membrane phospholipid composition and drives lipid mediator production via COX and LOX (Fig. 8I). Our data further suggest that LDs control canonical, PLA_2_-induced lipid mediator biosynthesis pathways by sequestering exogenous and membrane-derived PUFAs via DGAT1 and by modulating membrane PUFA content by delivering PUFAs into phospholipids via ATGL. We demonstrate that lipid mediator production induced by hGX sPLA_2_, which initially releases PUFAs from the plasma membrane and serum lipoproteins, depends on DGAT-driven enrichment of TAGs with PUFAs as well as ATGL-mediated lipolysis. Lipid mediator production induced by cPLA_2_α, which acts on perinuclear membranes to release AA for eicosanoid production, also depends on intact TAG turnover. These findings change the paradigm of PLA_2_-mediated inflammatory and mitogenic signalling by integrating membrane hydrolysis with LD metabolism. Furthermore, inhibition of DGAT-mediated TAG synthesis compromises lipid mediator release from cancer cells, thereby suppressing basal, sPLA_2_- and cPLA_2_α-induced cancer cell proliferation and reducing tumour growth in vivo. This study identifies the LD organelle as a central lipid trafficking hub that controls major PUFA supply routes for lipid mediator biosynthesis.

One of the most striking findings of the present study is that LD turnover regulates eicosanoid production from both exogenously-added PUFAs and membrane-hydrolysis-derived PUFAs suggesting that the build-up of PUFA-rich TAG stores is a required step in the control of lipid mediator signalling. A similar mechanism has been described in cardiomyocytes, whereby exogenous FAs have to be esterified into TAGs and then released by ATGL to activate peroxisome-proliferator-activated receptor (PPAR) signalling (Haemmerle *et al*, 2011; Zechner *et al*, 2012). Furthermore, as we found DGAT activity to be a prerequisite for both ATGL-induced and sPLA_2_/cPLA_2_α-induced eicosanoid production, it can be assumed that LDs control PUFA availability for eicosanoid production through the regulation of both TAG and membrane phospholipid pools.

Our previous data showed that breast cancer cells challenged with exogenous PUFAs depend on the balance between DGAT1-mediated sequestration of PUFAs into LDs and their release via ATGL-mediated lipolysis to survive lethal oxidative damage (Jarc *et al*, 2018). In support of this, inhibition of DGAT activity diverts dietary PUFAs towards esterification into membrane phospholipids, thereby increasing their peroxidation and leading to ferroptosis in acidic tumours (Dierge *et al*, 2021). Accordingly, DGAT1 increases the resilience of cancer cells against the stress of increased FA acquisition, which is a hallmark of transformed cells (Wilcock *et al*, 2022). It was recently shown that DGAT1-mediated TAG synthesis is required for prostaglandin formation during macrophage activation (Castoldi *et al*, 2020) and *Drosophila* oogenesis (preprint: Giedt *et al*, 2021). Our results suggest that DGAT activity controls cancer cell growth and proliferation by generating PUFA stores in LDs, which can be used for the production of potent, albeit short-lived, pro-inflammatory and anti-inflammatory lipid signalling molecules. As these signalling molecules are released from cells to act in autocrine and paracrine manners, they can modulate tumour growth by altering the function of cancer cells, immune cells and other cells in the tumour microenvironment (Wang & DuBois, 2010; Greene *et al*, 2011). Thus, in addition to their protective role against nutrient deficiency, lipotoxicity and oxidative stress (Petan, 2020), LDs modulate tumour growth also through the control of lipid mediator-dependent cell-autonomous and non-cell-autonomous mitogenic and inflammatory signalling pathways, and thus emerge as targets for therapeutic intervention. Targeting the DGAT enzymes might improve cancer treatments, particularly under conditions of elevated lipid influx (e.g., abundance of dietary fats, dyslipidemia, oncogene-driven elevated endogenous FA synthesis, high autophagic flux) (Chitraju *et al*, 2017; Nguyen *et al*, 2017; Dierge *et al*, 2021).

sPLA_2_ enzymes display tissue- and cell-specific expression patterns and differ in their membrane binding affinities and hydrolytic activities depending on the phospholipid composition and structural features of their target lipid assemblies (Lambeau & Gelb, 2008; Murakami *et al*, 2020). Their ability to promote lipid mediator production has been attributed either to direct PUFA release from phospholipids (e.g., in the case of hGX sPLA_2_) or cooperative action with cPLA_2_α (e.g., in the case of the group IIA sPLA_2_) (Hanasaki *et al*, 1999; Bezzine *et al*, 2002; Saiga *et al*, 2005). Regardless of the particular PLA_2_(s) involved, the release of PUFAs from phospholipids has been considered the final step in the control of PUFA entry into oxygenation pathways. Here, we show that lipid mediator production by two of the most potent mammalian PLA_2_s, hGX sPLA_2_ and cPLA_2_α, depends on LD turnover. Notably, hGX-sPLA_2_-treated MDA-MB-231 breast cancer cells showed an enrichment of both TAGs and phospholipids with PUFAs, but their release from TAGs by ATGL was essential for their conversion into lipid mediators and preceded the action of cPLA_2_α. Intriguingly, in A549 lung cancer cells, cPLA_2_α rather than ATGL was required for hGX-sPLA_2_-induced PGE_2_ production, but intact TAG synthesis was still a prerequisite for both hGX-sPLA_2_-induced and cPLA_2_α-induced eicosanoid synthesis and cell proliferation. This suggests that lipid mediator production could be mediated in different cell types by various combinations of PLA_2_s and (TAG) lipases, but the LD emerges as a central hub that controls the trafficking and final destination of PUFAs.

This study provides a novel view of PLA_2_-mediated inflammatory and mitogenic signalling, which integrates membrane hydrolysis with cellular LD metabolism. LDs could be the missing link that will help explain the elusive cross-talk between sPLA_2_s and cPLA_2_α in lipid mediator biosynthesis (Mounier *et al*, 2004; Saiga *et al*, 2005; Lambeau and Gelb, 2008). It will be interesting to investigate in future studies whether LD metabolism governs PLA_2_-induced lipid mediator production in various pathophysiological settings. Finally, the hGX sPLA_2_-induced incorporation of ω-3 and ω-6 PUFAs into TAGs, which is a novel mechanism of enzymatically-induced enrichment of LDs with phospholipid-derived PUFAs, could help clarify the various proposed roles of the group X sPLA_2_ that extend beyond lipid mediator production (Li *et al*, 2010; Sato *et al*, 2011; Murase *et al*, 2016; Schewe *et al*, 2016; Murakami *et al*, 2020). For example, PUFA-enriched TAGs have been observed in visceral adipose tissue of patients with colorectal cancer, along with elevated expression of hGX sPLA_2_ and prostaglandin biosynthetic enzymes (Liesenfeld *et al*, 2015).

Our results indicate that different cell-type-specific mechanisms might explain the dependence of cPLA_2_α-induced eicosanoid production on LDs. In the MDA-MB-231 and HeLa cells, cPLA_2_α depends on TAG synthesis and ATGL-mediated TAG lipolysis to drive the incorporation of PUFAs into phospholipids, which are then targeted by cPLA_2_α. In A549 cells, cPLA_2_α controls basal and sPLA_2_-induced PGE_2_ production independently of ATGL, although its activity still depends on intact LD biogenesis. In both cases, our data suggest that cPLA_2_α-mediated AA release for eicosanoid production occurs downstream of, and is controlled by, LD turnover. Interestingly, cPLA_2_α has also been shown to drive mitochondrial β-oxidation of both FAs and eicosanoids for energy production (Slatter *et al*, 2016), which is another indication that ATGL and cPLA_2_α share common metabolic and signalling pathways. Altogether, our findings suggest that LD turnover regulates the supply of PUFAs for lipid mediator production via at least two pathways: direct delivery of LD-derived PUFAs, which is independent of cPLA_2_α; and an indirect route, which requires cPLA_2_α but depends on LDs for the control of PUFA availability in membrane phospholipids. Notably, our data indicate that other lipases and LD breakdown mechanisms (e.g., lipophagy) besides (or instead of) ATGL might contribute to LD-dependent lipid mediator biosynthesis.

One of the intriguing questions that remains to be addressed in future studies is whether PUFA trafficking between the TAG core and the LD phospholipid monolayer is relevant for lipid mediator production. In principle, TAG lipolysis-derived PUFAs can be re-esterified in monolayer phospholipids and targeted by cPLA_2_α or other PLA_2_s. In agreement with this, cPLA_2_α, COXs, several prostaglandin synthetases and *de novo* produced eicosanoids have been localised to LDs (Wooten *et al*, 2008; Moreira *et al*, 2009; Bozza *et al*, 2011; Cruz *et al*,2020; Ward *et al*, 2020). In addition, cPLA_2_α is involved in remodelling of membrane shape and it might affect LD biogenesis and lipolysis by altering membrane composition and its biophysical properties (Gubern *et al*, 2008; Guijas *et al*, 2014; Antonny *et al*, 2015; Astudillo *et al*, 2019; Cao *et al*, 2019; Jarc & Petan, 2020, Zoni *et al*, 2021). In agreement with this, our results indicate that cPLA_2_α modulates membrane PUFA-phospholipid content and LD metabolism, including LD abundance and TAG acyl-chain composition. The molecular basis and functional relevance of these findings are currently unclear. We speculate that eicosanoid production does not necessarily occur on LDs as isolated cytosolic platforms, but at specific LD–ER contact sites that enable rapid lipid and protein transfer between the structures involved (i.e., the ER membrane bilayer, the LD phospholipid monolayer and TAG core) (Schuldiner & Bohnert, 2017). Such compartmentalisation would support the interplay among cPLA_2_α, ATGL and other enzymes in the control of LD turnover, membrane remodelling and the eicosanoid synthesis machinery.

In conclusion, our study reveals that LDs are essential for lipid mediator production and cancer cell proliferation. Esterification of PUFAs into TAGs by the DGAT enzymes and PUFA release from LDs (which depends on TAG lipolysis by ATGL, and/or other lipases depending on the cell type) drives lipid mediator production either directly, by feeding PUFAs into the COX/LOX machinery, or by redirecting some of the PUFAs into membrane phospholipids first, whereby they are then targeted by cPLA_2_α. Notably, both exogenous and sPLA_2_-membrane-hydrolysis-derived PUFAs have to cycle through LDs to be converted into lipid mediators, which reveals that LDs are optimal storage reservoirs for excess PUFAs that actively control PUFA release. LDs thus have an important regulatory role in lipid mediator production, thereby potentially affecting numerous downstream signalling pathways that are involved in inflammation, immunity and cancer.

## Materials and methods

### Materials

MDA-MB-231 human breast adenocarcinoma cells, A549 human lung carcinoma cells, HeLa human cervical adenocarcinoma cells, C2C12 mouse myoblasts cells, and Caco-2 human colorectal adenocarcinoma cells were obtained from American Type Culture Collection (ATCC, USA). J774A.1 mouse reticulum cell sarcoma macrophages were from the European Collection of Authenticated Cell Cultures (ECACC, UK), and MDA-MB-231 human breast adenocarcinoma cells with stable luciferase 2A and RFP expression (MDA-MB-231/Luciferase-2A-RFP) were from GeneTarget (USA). PC-3 human prostate adenocarcinoma cells were a kind gift from Dr. Mojca Pavlin (University of Ljubljana, Slovenia). OV-90 human ovarian papillary serous adenocarcinoma cells, TOV-112D human ovarian endometrioid carcinoma cells, TOV-21G human ovarian clear cell carcinoma cells and Ishikawa human endometrial adenocarcinoma cells were a kind gift from Dr. Brett McKinnon (Berne University Hospital, Switzerland). SGBS human Simpson-Golabi-Behmel syndrome preadipocytes were a kind gift from Dr. Merce Miranda (Joan XXIII University Hospital Tarragona, Spain), and SH-SY5Y human neuroblastoma cells were a kind gift from Dr. Boris Rogelj (Jožef Stefan Institute, Slovenia). RPMI-1640 culture medium was from ATCC (USA), and Dulbecco’s modified Eagle’s medium nutrient mixture F-12 (DMEM/F12), DMEM with high glucose and GlutaMAX supplement (DMEM-GlutaMax), foetal bovine serum, Dulbecco’s phosphate-buffered saline (DPBS), TrypLE Select and Opti-MEM were from Life Technologies (USA). AA and PGE_2_ standards were from Cayman Chemical (USA), and BODIPY 493/503, Lipofectamine RNAiMAX, Lipofectamine 3000, High-capacity cDNA reverse transformation kits were from Thermo Fisher Scientific (USA). Hoechst 33342 nuclear stain was from Enzo Life Sciences (USA). Human ATGL-targeting and cPLA_2_α-targeting siRNAs and the AllStars Negative Control siRNA were from Qiagen (Germany). T863 (DGAT1 inhibitor), PF-06424439 (DGAT2 inhibitor), indomethacin (COX inhibitor), nordihydroguaiaretic acid (LOX inhibitor), essentially fatty acid-free (EFAF) bovine serum albumin (BSA) (cat. no. A7511), FAF-BSA (cat. no. A8806) and Nile red were from Sigma-Aldrich (USA). High Pure RNA isolation kits were from Roche (Germany), horseradish-peroxidase-labelled secondary antibodies were from Jackson ImmunoResearch Laboratories (USA), ATGL antibodies (cat. no. #2138) were from Cell Signaling Technology (USA), cPLA_2_α antibodies (cat. no. sc-454) were from Santa Cruz (USA) and β-actin antibodies (cat. no. NB600-532) were from Novus Biologicals (UK). Recombinant wild-type hGX sPLA_2_ was prepared as described previously (Pucer *et al*, 2013). The full-length cDNAs coding for human cPLA_2_α (NCBI RefSeq NM.024420.3) and ATGL (NCBI RefSeq NM.020376.4) were cloned into the pcDNA 4/HisMaxC vector (Thermo Fisher Scientific, USA) using Gibson assembly cloning kits (New England Biolabs, USA), after removal of the N-terminal His-tag region. All of the other chemicals were of at least analytical grade, and were purchased from Sigma-Aldrich (USA) or Serva (Germany).

### Cell culture and treatments

MDA-MB-231 cells were cultured in RPMI-1640 medium, A549 cells in DMEM/F12 medium containing 2 mM L-glutamine (Gibco, USA), and HeLa cells in DMEM-Glutamax medium, all supplemented with 10% foetal bovine serum. Adherent cells were detached using TrypLE Select. Unless otherwise indicated, the cells were seeded in 24-well plates at a density of 3 ×10^4^ (MDA-MB-231), 2.5 ×10^4^ (A549) or 1.5 ×10^4^ (HeLa) cells/well and grown for 48 h in complete medium, followed by 24 h of serum deprivation in their respective media containing 0.02% EFAF-BSA. Aliquots of stock solutions of AA and PGE_2_ in absolute ethanol were stored under argon at −80 °C. Prior to addition to cell cultures, AA was resuspended in the relevant complete medium and incubated for 1 h at room temperature. Unless otherwise indicated, T863 and PF-06424439 were added to cells 2 h before treatments with recombinant hGX sPLA_2_ (1–10 nM) or AA (10 μM), and were present in the medium for the duration of the treatments. Prior to addition to cell culture, radiolabelled [^14^C]-OA was saponified by removing ethanol from the stock aliquot of [^14^C]-OA and resuspending in 50 μL 0.1 mM NaOH. Saponified [^14^C]-OA was incubated in the relevant complete medium for 30 min at room temperature and stored at –20 °C.

### Silencing of ATGL and cPLA_2_α expression using small-interfering RNAs

Reverse transfection was performed in 24-well plates at cell densities of 6 ×10^4^ (MDA-MB-231), 5 ×10^4^ (A549) or 3 ×10^4^ (HeLa) cells/well, or in 6-well plates at 3 ×10^5^ (MDA-MB-231), 2.5 ×10^5^ (A549) or 1.5 ×10^5^ (HeLa) cells/well. Gene expression silencing was performed with a 20 nM mixture of two ATGL-specific siRNAs (10 nM each) or a 40 nM mixture of four siRNAs specific for cPLA_2_α. Non-targeting siRNA controls contained 20 nM (for ATGL) or 40 nM (for cPLA_2_α) AllStars Negative Control siRNA (Qiagen). Transfection complexes were generated using 1 μL/well Lipofectamine RNAiMAX in 24-well plates, or 7.5 μL/well in 6-well plates, with Opti-MEM medium, according to manufacturer instructions.

### Transient overexpression of ATGL and cPLA_2_α

For transient overexpression experiments, the cells were seeded in complete medium in 24-well plates at a density of 9 ×10^4^ (MDA-MB-231), 6 ×10^4^ (A549) and 4.5 ×10^4^ (HeLa) cells/well, or in 6-well plates at 4.5 ×10^5^ (MDA-MB-231), 3 ×10^5^ (A549) and 2.5 ×10^5^ (HeLa) cells/well. Cells were then incubated for 24 h in complete medium, washed, and transfected with 0.5 μg/well plasmid DNA in 24-well plates, or 2.5 μg/well in 6-well plates, using Lipofectamine 3000 and Opti-MEM medium, according to manufacturer instructions; they were then left for 6 h in serum-depleted medium containing 0.02% EFAF-BSA (serum starvation). After 6 h, the cells were washed and treated according to the experimental set-up.

### Real-time quantitative PCR

Real-time quantitative (q)PCR analysis was performed as described previously (Pucer *et al*,2013; Brglez *et al*, 2014b). Briefly, the cells were seeded in complete medium in 6-well plates at 3 ×10^5^ (MDA-MB-231), 2.5 ×10^5^ (A549) and 1.5 ×10^5^ (HeLa cells) cells/well, grown 48 h in complete medium, followed by 24 h under serum deprivation in medium containing 0.02% EFAF-BSA. Total RNA was isolated from cell lysates and first-strand cDNA was generated using High-Capacity cDNA Reverse Transcription kits (Applied Biosystems, USA), according to the manufacturer instructions. qPCR analysis was performed on a StepOnePlus real-time PCR system (Thermo Scientific, USA) using FastStart Universal SYBR Green Master (Rox; Roche, Switzerland). Calibrator cDNA was transcribed from Quantitative PCR Human Reference Total RNA (Agilent Technologies, USA). Relative gene expression was calculated upon normalisation to two reference genes, considering primer-specific PCR efficiency and error propagation.

### Western blotting

The cells were seeded in complete medium in 6-well plates and reverse transfected with siRNAs and/or transiently transfected with pDNA, as described above. Cell lysates were prepared by scrapping adherent cells in Tris-glycine sodium dodecyl sulphate (SDS) sample buffer (2×) (Novex, Life Technologies, USA) that contained 800 mM dithiothreitol (Sigma-Aldrich, USA), with the addition of Halt protease inhibitor cocktail (Thermo Scientific, USA). Lysates were incubated at 95 °C for 10 min and stored on ice. Total protein concentrations were determined using Pierce 660 nm protein assays (Thermo Scientific, USA). Proteins (10–40 μg) were separated on 10% SDS-PAGE gels and then transferred to nitrocellulose membranes (Serva, Germany). The membranes were blocked for 1 h (for ATGL) or 2 h (for cPLA_2_α) in 5% non-fat dry milk in TBS/0.1% Tween-20 (TBST) or in 1% Western blocking reagent (WBR) (Roche Applied Science, Germany) in TBS (for β-actin), and incubated overnight at 4 °C in the presence of rabbit anti-human primary antibodies for ATGL (1:1000 dilution) or mouse anti-human primary antibodies for cPLA_2_α (1:250 dilution), both in 5% non-fat dry milk in TBST, or in rabbit anti-human primary antibodies for β-actin (1:5000 dilution) in 0.5% WBR in TBS. After washing with TBST, the membranes were incubated for 1 h with horseradish-peroxidase-conjugated secondary antibodies (1:10,000 dilution) in 5% non-fat dry milk in TBST for ATGL, in 0.5% WBR in TBST for cPLA_2_α, and in 0.5% WBR in TBS for β-actin. The signals were visualised using Lumi-Light Western Blotting Substrate (Roche Applied Science, Germany) on a Gel Doc XR system (Bio-Rad, USA).

### Neutral lipid quantification by flow cytometry

Cellular neutral lipid levels were quantified by flow cytometry as described previously (Pucer *et al*, 2013). Floating and adherent cells were harvested and centrifuged at 300× *g* for 10 min, and the pellets were resuspended in 500 μL 1 μg/mL Nile Red solution in DPBS. After a 10-min incubation in the dark, cell analysis was performed by flow cytometry on a FACSCalibur system, equipped with a 488-nm Ar-ion laser, and using the CellQuest software (Becton Dickinson, USA) and an FL-1 (530/30) filter, for at least 2 ×10^4^ events per sample.

### Triglyceride assays

Cellular TAG contents were determined using TAG assay kits - quantification (Abcam, USA). MDA-MB-231 and HeLa cells were seeded in 6-well plates at 1.5 ×10^5^ and 7 ×10^4^ cells/well, respectively. After 24 h, MDA-MB-231 cells were treated with 10 nM hGX sPLA_2_ and HeLa cells with 1 nM hGX sPLA_2_ in complete medium for 48 h. Cell lysates were prepared and used for TAG quantification according to the manufacturer instructions.

### Glycerol release assays

Cellular lipolytic activity was assessed by measuring glycerol release in cell supernatants. Briefly, the cells were reverse transfected and seeded in 48-well plates at a density of 3 ×10^4^ (MDA-MB-231 cells) or 1.5 ×10^4^ cells/well (HeLa cells). For reverse transfection, 0.5 μL Lipofectamine RNAiMAX, 20 nM siRNA and 40 μL OPTI-MEM medium were used per well. After 24 h, the cells were washed, placed in serum-starvation medium containing 0.02% EFAF-BSA, and transfected with 0.250 μg plasmid DNA for protein overexpression using Lipofectamine 3000, according to the manufacturer instructions. After 6 h, the cells were washed and treated with 10 nM hGX sPLA_2_ for 48 h in complete medium. Cell supernatants were collected in low-binding microcentrifuge tubes and centrifuged for 10 min (4 °C, 16,000× *g*), and the glycerol concentrations were determined using the Glycerol Cell-Based assay kits (Cayman Chemicals, USA), according to the manufacturer instructions.

### Untargeted lipidomic analysis of phospholipids and triglycerides

For untargeted lipidomic analysis of hGX sPLA_2_-induced changes in TAG acyl-chain composition, MDA-MB-231 cells were seeded in complete medium on 10-cm plates at 1 ×10^6^ cells/plate. After 24 h, the cells were treated with 1 nM hGX sPLA_2_ in complete medium for 48 h. Cell lysates were prepared by washing the cells twice with DPBS and scraping in 1 mL lysis buffer (20 mM Tris-HCl, pH 7.4, 2 mM EDTA, 2 μL Halt protease inhibitor cocktail), followed by centrifugation for 10 min (1000× *g*, 4 °C). Cell pellets were resuspended in 150 μL lysis buffer and sonicated on ice. Total lipids were extracted in chloroform/methanol (2/1, v/v) containing 1% acetic acid, 500 nM butylated hydroxytoluene (BHT) and internal standards (IS; 100 pmol 17:0/17:0/17:0 triacylglycerol, Larodan, Solna, Sweden) under constant shaking for 1 h (30 rpm/min, 4 °C). After centrifugation at 3300 rpm for 20 min at room temperature, the upper aqueous layer was removed, and the organic solvents were evaporated using a sample concentrator (Techne, UK) equipped with the Dri-Block DB-3 heater (Techne, UK). Lipids were resolved in 200 μL chloroform and stored at −20 °C. Prior to mass spectrometry, the samples were placed at room temperature and dried and resuspended in 1 mL chloroform/methanol (2/1, v/v). An aliquot of each sample (20 μL) was mixed with 180 μL isopropanol, and 5 μL was used for chromatographic separation on an Acquity-UPLC system (Waters Corporation, Milford, MA, USA), equipped with an ACQUITY BEH C18 column (2.1 × 50 mm, 1.7 μm; Waters Corporation, Milford, MA, USA). A SYNAPT_™_G1 qTOF HD mass spectrometer (Waters Corporation, Milford, MA, USA) equipped with an ESI source was used for detection. Data acquisition was carried out using the MassLynx 4.1 software (Waters), and the lipid classes were analysed with the Lipid Data Analyser 1.6.2 software. The data were normalised for recovery, extraction and ionisation efficacy by calculating analyte/internal standard ratios (AU) and expressed as percentage composition.

To determine the changes in the phospholipid and TAG profiles in ATGL-depleted and cPLA_2_α-depleted cells, MDA-MB-231 cells were seeded in complete medium on 6-well plates at 3 ×10^5^ cells/well and reverse transfected with ATGL-specific and/or cPLA_2_α-specific siRNAs, as described above. After 24 h, the cells were washed and grown for 48 h in complete medium in the presence or absence of 10 nM sPLA_2_. The cells were then washed twice with DPBS and serum starved for the following 24 h in RPMI-1640 medium containing 0.02% EFAF-BSA. Samples were collected (at 0, 3, 24 h of serum starvation) by placing the plates on ice, washing the cells twice with ice-cold DPBS, and scraping the cells in 300 μL lysis buffer, followed by centrifugation for 10 min (1000× *g*, 4 °C). The cell pellets were resuspended in 150 μL lysis buffer and sonicated on ice. Then, 10 μL of each sample was used for protein determination (Pierce 660). Total lipids were extracted by transferring the pellets into 2 ml tubes followed by homogenisation in 700 μL of a 3:1 (v/v) mixture of methyl tert-butyl ether and methanol, containing 1% acetic acid, 500 nM BHT and IS (8 pmol 18:3/18:3/18:3 triacylglycerol, 14:0/14:0 phosphatidylcholine, Larodan, Solna, Sweden; 50 pmol 17:0/17:0 phosphatidylethanolamine, 12 pmol 17:0/17:0 phosphatidylserine, Avanti Polar Lipids, Alabaster, AL, USA), with two steel beads on a mixer mill (30 Hz, Retsch, Germany) at 4 °C. After homogenisation, the samples were mixed at room temperature under constant shaking for 30 min. Then 140 μL distilled H_2_O was added, and the samples were thoroughly mixed and centrifugated (14,000 rpm, 10 min), to establish phase separation. The organic phase (500 μL) was transferred into new tubes and the organic solvent was evaporated off under a stream of nitrogen. The residual protein slurry was dried and used for protein determination after lysis in 400 μL NaOH/SDS (0.3 M/0.1%). Prior to mass spectrometry analysis, the lipids were resolved in 200 μL isopropanol /methanol/H_2_O (70/25/10, v/v/v). Chromatographic separation was performed on a 1290 Infinity II LC system (Agilent, Santa Clara, CA, USA) equipped with a C18 column (Zorbax RRHD Extend; 2.1 × 50 mm, 1.8 μm; Agilent, Santa Clara, CA, USA), using a 16 min linear gradient from 60% solvent A (H_2_O; 10 mM ammonium acetate, 0.1% formic acid, 8 μM phosphoric acid) to 100% solvent B (2-propanol; 10 mM ammonium acetate, 0.1% formic acid, 8 μM phosphoric acid). The column compartment was kept at 50 °C. A Q-TOF mass spectrometer (6560 Ion Mobility; Agilent, Santa Clara, CA, USA) equipped with electrospray ionisation source (Dual AJS) was used for detection of the lipids in positive and negative Q-TOF mode. Data acquisition was carried out using the MassHunter Data Acquisition software (B.09; Agilent, Santa Clara, CA, USA). Lipid species were manually identified and lipid data were processed using MassHunter Quantitative Analysis (B.09.00; Agilent, Santa Clara, CA, USA). Data were normalised for recovery, extraction and ionisation efficiency by calculation of the analyte/ internal standard ratios, and are expressed as fmol/μg protein. Lipidomic data were analysed by multiple t-test analysis of log-transformed data to compare two conditions at a time, using GraphPad Prism 9.0.2 (GraphPad Software, USA). Lipid ontology enrichment analysis was performed using the LION/web enrichment and principal component analysis modules (Molenaar *et al*, 2019).

### Thin layer chromatography of radiolabelled cellular lipids

MDA-MB-231 cells were seeded in complete medium on T-25 flasks at a density of 5 ×10^5^ cells/flask. After 24 h, the cells were treated with 1 μCi/sample [^14^C]-OA for 18 h, washed twice with DPBS, and treated with 10 nM hGX sPLA_2_ under three different conditions: (a) 24 h in complete medium; (b) 24 h in complete medium followed by 96 h serum deprivation in 0.02% EFAF-BSA in RPMI-1640; and (c) 24 h in 0.02% FAF-BSA in RPMI-1640 followed by 96 h in 0.02% EFAF-BSA in RPMI-1640. Lipid extraction was performed with 1 mL hexane/ isopropanol (3:2, v/v) under constant shaking for 10 min at room temperature, and repeated twice. The samples were stored in microcentrifuge tubes at −20 °C. Total proteins were isolated from cell remnants in 2 mL lysis buffer (0.3 M NaOH, 0.1% SDS) under constant shaking for 2 h at room temperature. Protein concentrations were determined using BSA standard solutions (Thermo Scientific, USA) and the BCA protein assay reagent (Thermo Scientific, USA). Samples of cellular lipids were dried, resuspended in 20 μL chloroform (repeated three times), and loaded onto the stationary phase of silica TLC plates immobilised on a polymeric binder (Merck, Germany). Dry TLC plates were developed using chloroform/ methanol/ acetone/ acetic acid/ H_2_O (50/10/20/12/5, v/v/v/v/v) for phospholipid separation, and hexane/ diethyl ether/ acetic acid (70:29:1, v/v/v) for neutral lipid separation. Radiolabelled lipids were detected with autoradiography using an imager (PhosphorImager SI; Amersham Biosciences, UK). To quantify the TAG content, the TLC plates were incubated in an iodine steam, the TAG patches were cut out and placed in vials with scintillation cocktail, and the radioactivity was measured using a liquid scintillation counter (Tri-Carb 1600CA; PerkinElmer, USA).

### Lipid mediator analysis using UPLC-MS-MS

MDA-MB-231 cells were seeded in complete medium in 6-well plates at 3 × 10^5^ cells/well. In case of ATGL silencing, cells were reverse transfected with ATGL-specific siRNAs as described above. After 24 h, the cells were treated with 10 nM hGX sPLA_2_ and/or a mixture of 20 μM DGAT1 and 20 μM DGAT2 inhibitors for 48 h, washed with DPBS and left for 24 h in RPMI-1640 medium containing 0.02% EFAF-BSA. Cells with DGAT inhibition were treated with the inhibitors also during the serum starvation phase. The cells were lysed and total protein concentrations were determined as described above. Supernatants were collected in low-binding microcentrifuge tubes, centrifuged at 4 °C (16,000× *g*, 10 min) and stored at −80 °C before shipping on dry ice. 1 mL of the supernatants (MDA-MB-231 experiment) were first mixed with 2 mL of ice-cold methanol containing deuterium-labelled internal standards (200 nM d8-5S-HETE, d4-LTB_4_, d5-LXA_4_, d5-RvD2, d4-PGE_2_ and 10 μM d8-AA; Cayman Chemical/Biomol GmbH, Hamburg, Germany) to facilitate quantification and sample recovery. Samples were kept at −20 °C for 60 min to allow protein precipitation.

Tumour samples for UPLC-MS-MS analysis were collected from mice xenografts. After cancer cell injection and visible tumour growth, mice were subcutaneously treated daily with T863 at a dose of 9.6 mg/kg or corresponding vehicle (0.2% DMSO) for two weeks before being sacrificed and sampled for tumours. Tumour samples (30–50 mg) were weighed and 25 μL methanol of each mg tumour was added and then homogenized. After that, samples were centrifuged (16,000× g, 4 °C, 5 min), and 800 μL (corresponding to 32 mg tumour) were filled up to 2 mL methanol and 1 mL PBS was added containing deuterium-labelled internal standards (200 nM d8-5S-HETE, d4-LTB_4_, d5-LXA_4_, d5-RvD2, d4-PGE_2_ and 10 μM d8-AA; Cayman Chemical/Biomol GmbH) to facilitate quantification and sample recovery. Lipid mediator analysis using UPLC-MS-MS was performed as described previously (Werz *et al*, 2018) with some minor modifications (Werner *et al*, 2019). Briefly, after centrifugation (1200× *g*, 4 °C, 10 min) 8 mL of acidified H_2_O was added (final pH = 3.5) and the samples were subjected to solid phase extraction. The solid phase cartridges (Sep-Pak®Vac 6cc 500 mg/ 6 mL C18; Waters, Milford, MA) were equilibrated with 6 mL methanol and then with 2 mL H_2_O prior sample loading onto the columns. After washing with 6 mL H_2_O and additional 6 mL *n*-hexane, lipid mediators were eluted with 6 mL methyl formate. The eluates were brought to dryness using a TurboVap LV evaporation system (Biotage, Uppsala, Sweden) and resuspended in 200 μL methanol/water (50/50, v/v) for UPLC-MS-MS analysis. Lipid mediator profiling was analyzed with an Acquity^™^ UPLC system (Waters, Milford, MA, USA) and a QTRAP 5500 Mass Spectrometer (ABSciex, Darmstadt, Germany), equipped with a Turbo V™ Source and electrospray ionization. Lipid mediators were separated using an ACQUITY UPLC^®^BEH C18 column (1.7 μm, 2.1 × 100 mm; Waters, Eschborn, Germany) at 50 °C with a flow rate of 0.3 mL/min and a mobile phase consisting of methanol/water/acetic acid of 42/58/0.01 (v/v/v) that was ramped to 86/14/0.01 (v/v/v) over 12.5 min and then to 98/2/0.01 (v/v/v) for 3 min (Werner *et al*, 2019). The QTrap 5500 was operated in negative ionization mode using scheduled multiple reaction monitoring (MRM) coupled with information-dependent acquisition. The scheduled MRM window was 60 sec, optimized lipid mediator parameters were adopted (Werner *et al*, 2019), and the curtain gas pressure was set to 35 psi. The retention time and at least six diagnostic ions for each lipid mediator were confirmed by means of external standards (Cayman Chemical/Biomol GmbH). Quantification was achieved by calibration curves for each lipid mediator. Linear calibration curves were obtained for each lipid mediator and gave r^2^ values of 0.998 or higher (for fatty acids 0.95 or higher). Additionally, the limit of detection for each targeted lipid mediator was determined (Werner *et al*, 2019).

### Quantification of PGE_2_ in cell supernatants

The amounts of PGE_2_ released into culture medium were routinely determined using a commercial enzyme immunoassay (Prostaglandin E_2_ Express ELISA kits; Cayman Chemicals, USA). The cells were seeded in 24-well plates and reverse transfected with siRNAs as described above. After 24 h, the cells were transfected with plasmid DNA as described above and treated with 10 nM hGX sPLA_2_ and/or 10 μM AA and/or a mixture of 20 μM T863 and 20 μM PF-06424439 (DGAT inhibitors) for 48 h in complete medium. The cells were then washed and serum starved for 24 h in the appropriate culture medium containing 0.02% EFAF-BSA. In some experiments, DGAT inhibitors were included in the serum-starvation phase. Cell supernatants were collected in low-binding microcentrifuge tubes, centrifuged at 4 °C (16,000× *g*, 10 min) and immediately used for analysis or stored at −80 °C for up to 7 days. For analysis, 50 μL undiluted samples were used, and standard curves prepared by diluting the PGE_2_ standard in culture medium.

### Confocal microscopy

Cytosolic LDs were visualised using BODIPY 493/503 neutral lipid staining. Cells were reverse transfected and seeded on glass-bottomed culture plates at 6 ×10^4^ (MDA-MB-231 cells), 3 ×10^4^ (HeLa cells) and 5 ×10^4^ (A549 cells) cells/well. Twenty-four hours later, the media were replaced and the cells were treated with 10 nM hGX sPLA_2_ and the DGAT inhibitors in complete medium for 48 h, followed by 24-h starvation in serum-free medium containing 0.02% EFAF-BSA. For confocal microscopy, the cells were washed twice with DPBS, stained with 1 μg/mL BODIPY 493/503 and 1 μg/mL Hoechst stain solution in RPMI-1640 medium for 15 min in a CO_2_ incubator, washed twice with DPBS, and left in fresh RPMI-1640 medium. Live imaging was carried out at the beginning of and after the 24-h serum starvation using a confocal laser scanning microscope (LSM 710; Carl Zeiss, Germany) and a stage-top microscope CO_2_ incubation system (Tokai Hit, Japan). Images were processed using the Zen software (Carl Zeiss, Germany).

### LD counting and diameter analysis

Confocal microscopy images of BODIPY 493/503 stained LDs were used for computer image analysis with the ImageJ software (National Institutes of Health, USA) and the LD Counter Plugin (https://doi.org/10.5281/zenodo.2581434). Analysis was performed on 32-bit two-dimensional images, whereby the numbers and sizes of the LDs were determined according to the plugin instructions. LD diameters were calculated from the LD surface areas. Analyses were performed on at least 40 cells/sample. Data were loaded into the Prism 9.0.2 software (GraphPad Software, USA), with the geometric means of the LD diameters and numbers calculated per individual cell in the samples, followed by the statistical analysis.

### Cell proliferation

MDA-MB-231/Luciferase-2A-RFP cells were seeded in complete medium in 96-well plates at 5 ×10^3^ cells/well in at least triplicates. After 24 h, the cells were treated with 20 μM DGAT1 and 20 μM DGAT2 inhibitors and/or 1 μM PGE_2_ and/or 10 nM hGX sPLA_2_ as described above, and left in complete medium for 72 h. PGE_2_ was replenished every 24 h by direct addition to the culture medium. In transient transfection experiments, the cells were seeded in complete medium on 48-well plates at 3 ×10^4^ (MDA-MB-231/Luciferase-2A-RFP cells) or 1.5 ×10^4^ (A549 cells) cells/well. After 24 h, the cells were washed and transfected with 0.250 μg plasmid DNA/well using 0.5 μL P3000 reagent and 1 μL Lipofectamine 3000 in OptiMEM medium, and incubated in RPMI-1640 (MDA-MB-231) or DMEM/F12 with 2 mM L-glutamine (A549) containing 0.02% EFAF-BSA. After 6 h, the cells were treated with 10 nM recombinant hGX sPLA_2_ in complete medium, with 25 μM (A549) or 50 μM (MDA-MB-231) indomethacin, 25 μM nordihydroguaiaretic acid, and/or 20 μM each of the T863 and PF-06424439 DGAT1 and DGAT2 inhibitors, respectively. A549 cell proliferation was determined after 48 h using the direct cell proliferation assay kits (CyQUANT; Invitrogen, USA). MDA-MB-231/Luciferase-2A-RFP cell proliferation was determined by measuring RFP fluorescence emission (excitation, 558 nm; emission, 583 nm) on a microplate reader (Infinite M1000; Tecan, Austria). The final data were obtained after subtracting the background signal of the blank samples that contained culture medium without cells.

### Clonogenic assay

MDA-MB-231 cells were seeded in 24-well plates in complete medium as described above. After 24 h, the cells were treated with 20 μM T863 and/or 20 μM PF-06424439 and left for 24 h. Cells were then washed and harvested for cell counting. 500 cells/well were seeded in 6-well plates and colony formation followed for nine days in complete medium containing 20 μM T863 and/or 20 μM PF-06424439. Medium was replaced every 48–72 h. For analysis, colonies were washed with DPBS, fixed in 100% cold methanol at 4 °C for 15 min and stained with 0.5% crystal violet in DBPS. Images were taken for colony counting with ImageJ software (National Institutes of Health, USA).

### Mouse studies

All animal experiments were performed according to the directives of the EU 2010/63 and were approved by the Administration of the Republic of Slovenia for Food Safety, Veterinary and Plant Protection of the Ministry of Agriculture, Forestry and Foods, Republic of Slovenia (Permit Number U34401-9/2020/9). Laboratory animals were housed in IVC cages (Techniplast), fed standard chow (Mucedola) and tap water was provided ab libitum. Mice were maintained in 12-12 h dark-light cycle at approximately 40–60% relative humidity and an ambient temperature of 22 °C. All animals used in the study were healthy and accompanied with health certificate from the animal vendor. Health status was confirmed by FELASA recommended Mouse Vivum immunocompetent panel (QM Diagnostics). To test the anti-tumour effect of DGAT1 inhibition, female 8–10 weeks old SCID C.B-17/IcrHsd-*Prkdc^scid^* mice (Envigo, Italy) were used for xenograft cancer studies. 3×10^6^ MDA-MB-231/Luciferase-2A-RFP cells were implanted into the right flank of the mouse. When a certain tumour dimension reached 5 mm, subcutaneous daily treatments were started with T863 at a dose of 9.6 mg/kg or with corresponding 0.2% DMSO vehicle. Body weight and tumour size via calliper measurements were performed every couple of days. Tumour burden was calculated using the formula V=(a*b*c*Π)/6. Animals were humanely euthanized either when the observed tumour dimension reached 12 mm, the tumour ulcerated or when animals lost over 20% of their own body mass. Tumour samples at the end of the experiment were collected and snap frozen in liquid nitrogen for further UPLC-MS-MS analysis.

### Statistical analysis

Statistical analyses were performed using Prism 9.0.2 (GraphPad Software, USA). Unless otherwise indicated, the data are presented as means ±SEM of at least three independent experiments. Statistical significance was determined using t-tests, one-way or two-way ANOVA, followed by Bonferroni, Sidak or Tukey’s multiple comparison tests. P values <0.05 were considered as statistically significant.

## Acknowledgements

We are grateful to Ana Temprano Lopez, Petra Hruševar, Ana Kump, Ema Guštin and Belen Vilanova Baeza for their technical help, Dr. Mojca Pavlin (University of Ljubljana, Slovenia), Dr. Brett McKinnon (Berne University Hospital, Switzerland), Dr. Merce Miranda (Joan XXIII University Hospital Tarragona, Spain) and Dr. Boris Rogelj (Jožef Stefan Institute, Slovenia) for kindly providing cell lines. We thank Dr. Chris Berrie for critical reading of the manuscript. This work was supported by the Slovenian Research Agency young researcher (1000-15-106) and postdoctoral grants (Z3-2650) to E.J.J., the P1-0207 Programme grant, and J7-1818 Project grant to T.P., P4-0176 Programme grant to R.J. and J4-4563 Project grant to D.L.; by the Deutsche Forschungsgemeinschaft (DFG, German Research Foundation), Project-ID 239748522 - SFB 1127 ChemBioSys, and the EU COST Action CA19105 EpiLipidNet.

## Author contributions

E.J.J. performed most of the experiments, analysed the data, and prepared manuscript draft; A.P.J. and V.B. performed initial lipidomic experiments; Š.K. prepared samples for lipid mediator mass spectrometry analysis and performed LD diameter analysis; T.O.E. performed mass spectrometry analyses, designed experiments, contributed ideas and revised the paper; R.Z. provided materials, designed experiments, contributed ideas and revised the paper; G.L. contributed materials, ideas and revised the paper; P.M.J., J.G. and O.W. performed lipid mediator mass spectrometry analyses and revised the paper; D.L. and A.G.U. performed in vivo studies, and R.J. revised the paper; T.P. conceptualised the study, analysed the data, prepared the figures and illustrations, and wrote the manuscript.

## Supplementary Material

**Supplementary Figure 1.**
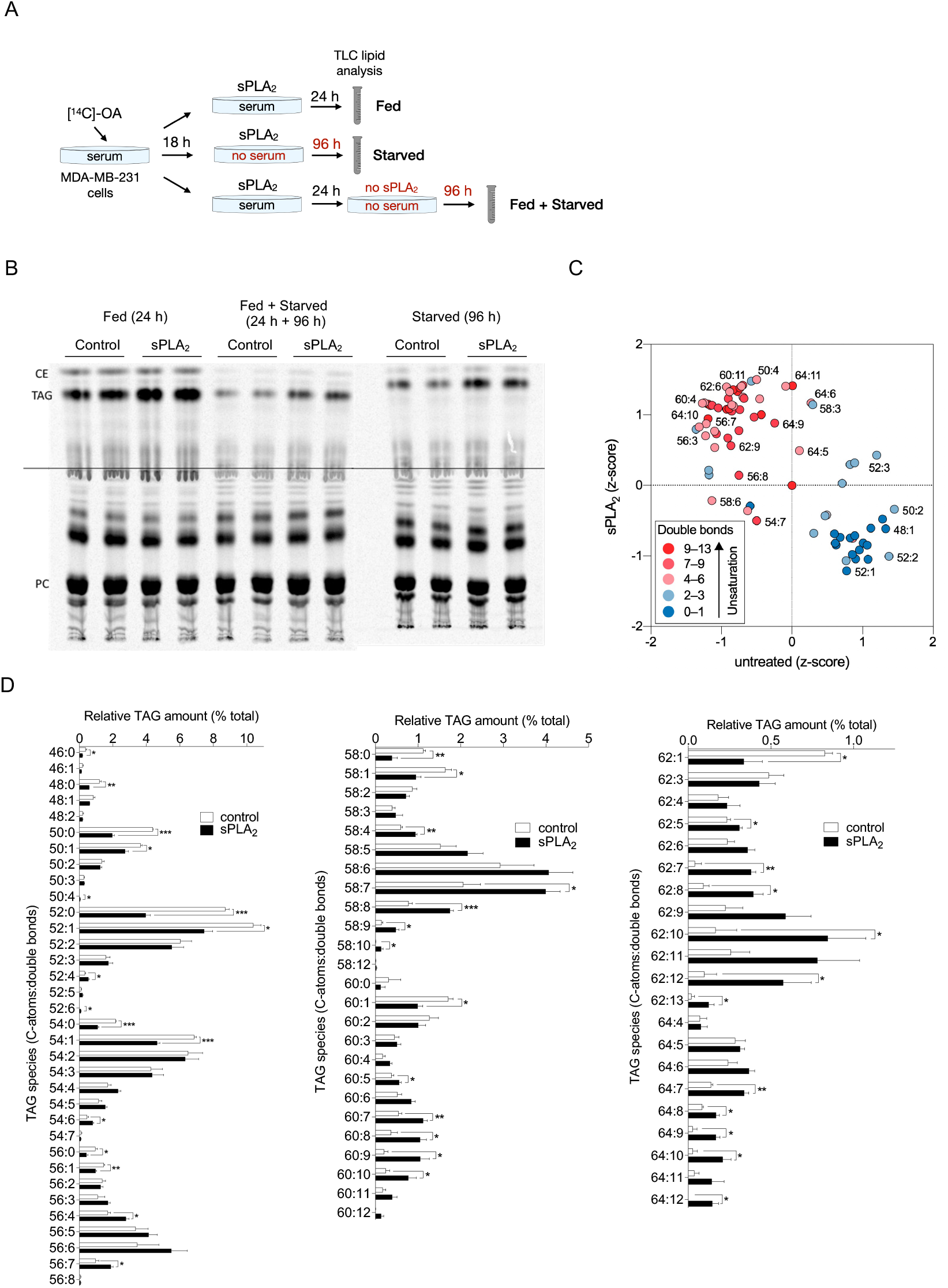
sPLA_2_-induced changes in LD metabolism and composition. (**A, B**) Diagram illustrating the experimental set-up for oleate incorporation analyses shown in (B) and in Figure 1D. (B) Representative thin layer chromatography (TLC) plate showing that in hGX-sPLA_2_-treated cells, oleate is preferentially incorporated into TAGs, but not into phosphatidylcholine (PC) or cholesterol esters (CE). (**C, D**) Lipidomic analysis of hGX-sPLA_2_-induced PUFA-TAG enrichment in MDA-MB-231 cells grown under serum-rich conditions. (C) Representative XY z-score plot showing colour-coded changes in the levels of TAG acyl-chain unsaturation. Data are means ±SEM of three or four (D) independent experiments. *, P <0.05; **, P <0.01; ***, P <0.001 (unpaired t-tests (D).

**Supplementary Figure 2.**
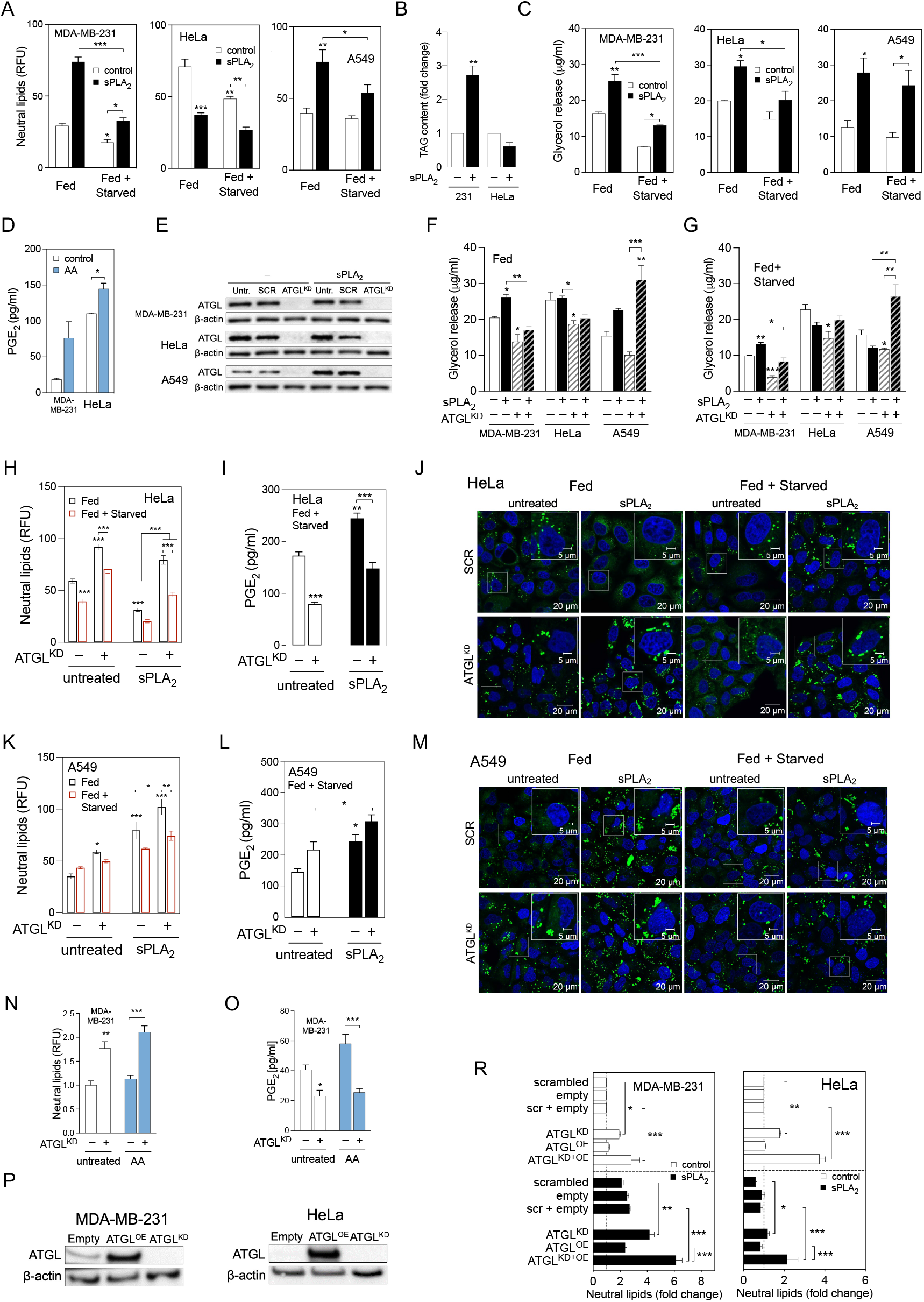
ATGL-mediated LD breakdown drives PGE_2_ production in starving MDA-MB-231 and HeLa cells, but not in A549 cells. (**A**) Neutral lipid contents in control and hGX-sPLA_2_-treated cells at the beginning of and after serum starvation. (**B**) Total cellular triglycerides (TAGs) in control and hGX-sPLA_2_-treated MDA-MB-231 and HeLa cells quantified using a biochemical assay. (**C**) Glycerol released from fed and starving control and hGX-sPLA_2_-treated cells. (**D**) PGE_2_ levels in cell supernatants of control and AA-treated cells. (**E**) Representative western blot showing siRNA-induced ATGL knock down (ATGL^KD^) in comparison with control (untransfected [untr.] and non-targeting siRNA-transfected [SCR] cells), treated and grown as shown in Fig. 2C. (**F, G**) Glycerol released in cell supernatants of control and ATGL-depleted serum-fed and serum-starved cells, either untreated or treated with hGX sPLA_2_. (**H, I, K, L**) Neutral lipid and PGE_2_ levels in ATGL-silenced control and sPLA_2_-treated cells grown and analysed as shown in Fig. 2C. (**J, M**) Representative confocal microscopy images showing effects of ATGL depletion on cellular LD content in control and hGX-sPLA_2_-treated cells, under serum-rich (Fed) and serum-free (Fed + Starved) conditions. LDs and nuclei were stained using BODIPY 493/503 and Hoechst 33342, respectively. (**N, O**) Changes in LD levels and PGE_2_ production induced by ATGL silencing in control and arachidonic acid (AA)-treated cells analysed as shown in Fig. 2C. (**P**) Representative western blot showing ATGL protein overexpression in cells transfected with ATGL-encoding plasmid (ATGL^OE^) in comparison with those transfected with control (Empty) vector or ATGL-specific siRNA (ATGL^KD^), grown as illustrated in Fig. 2G. (**R**) Neutral lipid levels in ATGL-overexpressing in untreated and hGX-sPLA_2_-pretreated serum-starved cells, in comparison with those co-transfected with ATGL-specific siRNA (ATGL^KD+OE^), non-targeting siRNA (scrambled) and control plasmid (empty), grown as shown in Fig. 2G. (A, H, K, N, R) Neutral lipids were quantified by Nile Red staining and flow cytometry. (D, I, L, O) PGE_2_ was quantified by ELISA at the end of starvation. Data are means ±SEM of two (D) or three (A–C, F–I, K, L, N, O) independent experiments. *, P <0.05; **, P <0.01; ***, P <0.001 (two-way ANOVA with Bonferroni (B, F–I, K, L, N, O) or Tukey (A, C, R) adjustment; unpaired t-test (D)).

**Supplementary Figure 3.**
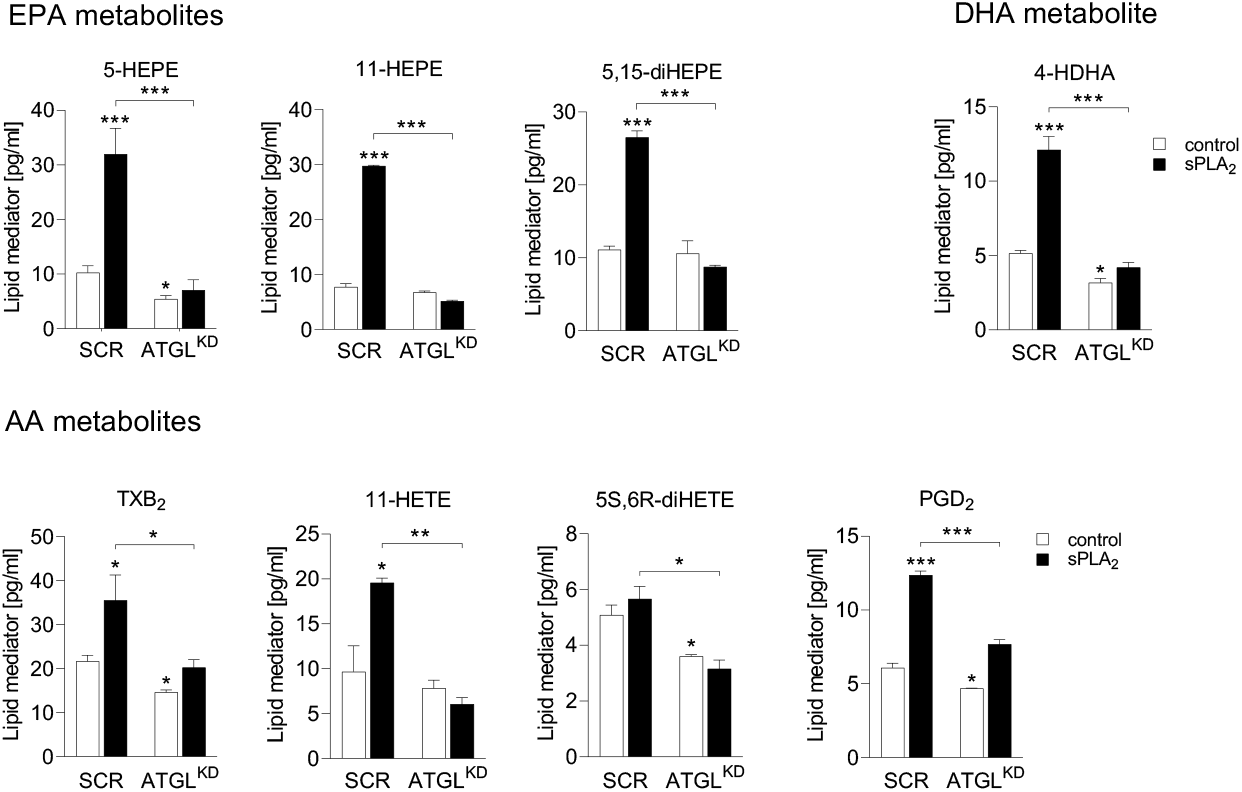
Depletion of ATGL suppresses both basal and hGX-sPLA_2_-stimulated lipid mediator production in breast cancer cells. hGX-sPLA_2_-induced and ATGL knockdown (ATGL^KD^)-induced changes in selected lipid mediators (as indicated) released from serum-starved MDA-MB-231 cells. Data are means ±SEM of three independent experiments. *, P <0.05; **, P <0.01; ***, P <0.001 (two-way ANOVA with Sidak adjustment). DHA, docosahexaenoic acid; 4-HDHA, 4-hydroxydocosahexaenoic acid; EPA, eicosapentaenoic acid; HEPE, hydroxyeicosapentaenoic acid; HETE, hydroxyeicosatetraenoic acid; AA, arachidonic acid; TXB_2_, thromboxane B_2_.

**Supplementary Figure 4.**
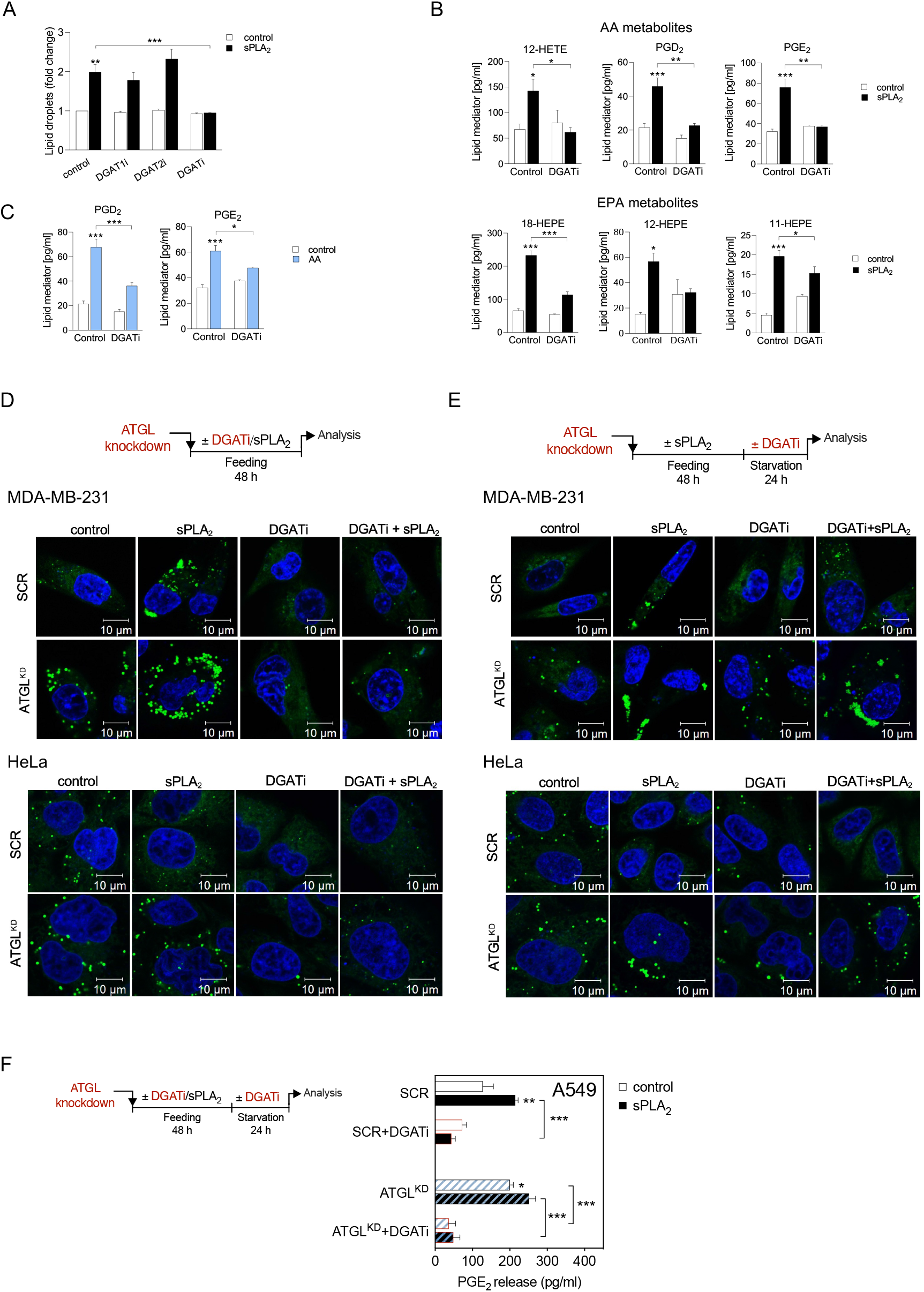
Lipid mediator production induced by hGX sPLA_2_ or exogenous AA depends on DGAT-mediated LD biogenesis. (**A**) Neutral lipid content in MDA-MB-231 cells grown in the absence and presence of T863 (DGAT1i) and PF-06427878 (DGAT2i), or an equimolar mix of both DGAT inhibitors (DGATi), without and with stimulation of LD biogenesis by hGX sPLA_2_ during serum sufficiency. (**B**) hGX-sPLA_2_-induced and DGAT inhibition (DGATi)-induced changes in selected lipid mediators (as indicated) released from serum-starved MDA-MB-231 cells. Cells were treated as shown in Figure 4A. (**C**) AA-induced and DGATi-induced changes in selected lipid mediators (as indicated). Cells were treated as shown in Figure 4A. (**D, E**) Diagrams illustrating the experimental conditions used (top) and representative live-cell confocal microscopy images of control (SCR) and ATGL-depleted (ATGL^KD^) MDA-MB-231 and HeLa cells treated with DGAT inhibitors during serum feeding (D) and during serum starvation (E), without and with hGX sPLA_2_ pre-treatment. LDs and nuclei were visualised using BODIPY 493/503 and Hoechst 33342 staining and confocal microscopy. (**F**) Diagram illustrating the experimental conditions (left) and graph showing DGAT inhibition (DGATi)-induced changes in PGE_2_ production in serum-starved control (SCR) and ATGL-depleted (ATGL^KD^) A549 cells, without and with stimulation of LD biogenesis by hGX sPLA_2_ pre-treatment. Neutral lipids were measured at the end of starvation with Nile Red staining and flow cytometry. Lipid mediators were quantified by UPLC-MS-MS. (F) PGE_2_ levels were determined in cell supernatants as described in Methods. Data are means ±SEM of three (A) or four (B, C) independent experiments. *, P <0.05; **, P <0.01; ***, P <0.001 (two-way ANOVA with Tukey (A), Sidak (B, C) or Bonferroni (F) adjustment). EPA, eicosapentaenoic acid; HEPE, hydroxyeicosapentaenoic acid; AA, arachidonic acid; TXB_2_, thromboxane B_2_; HDHA, hydroxydocosahexaenoic acid; HETE, hydroxyeicosatetraenoic acid; PGE_2_, prostaglandin E_2_; PGD_2_, prostaglandin D_2_.

**Supplementary Figure 5.**
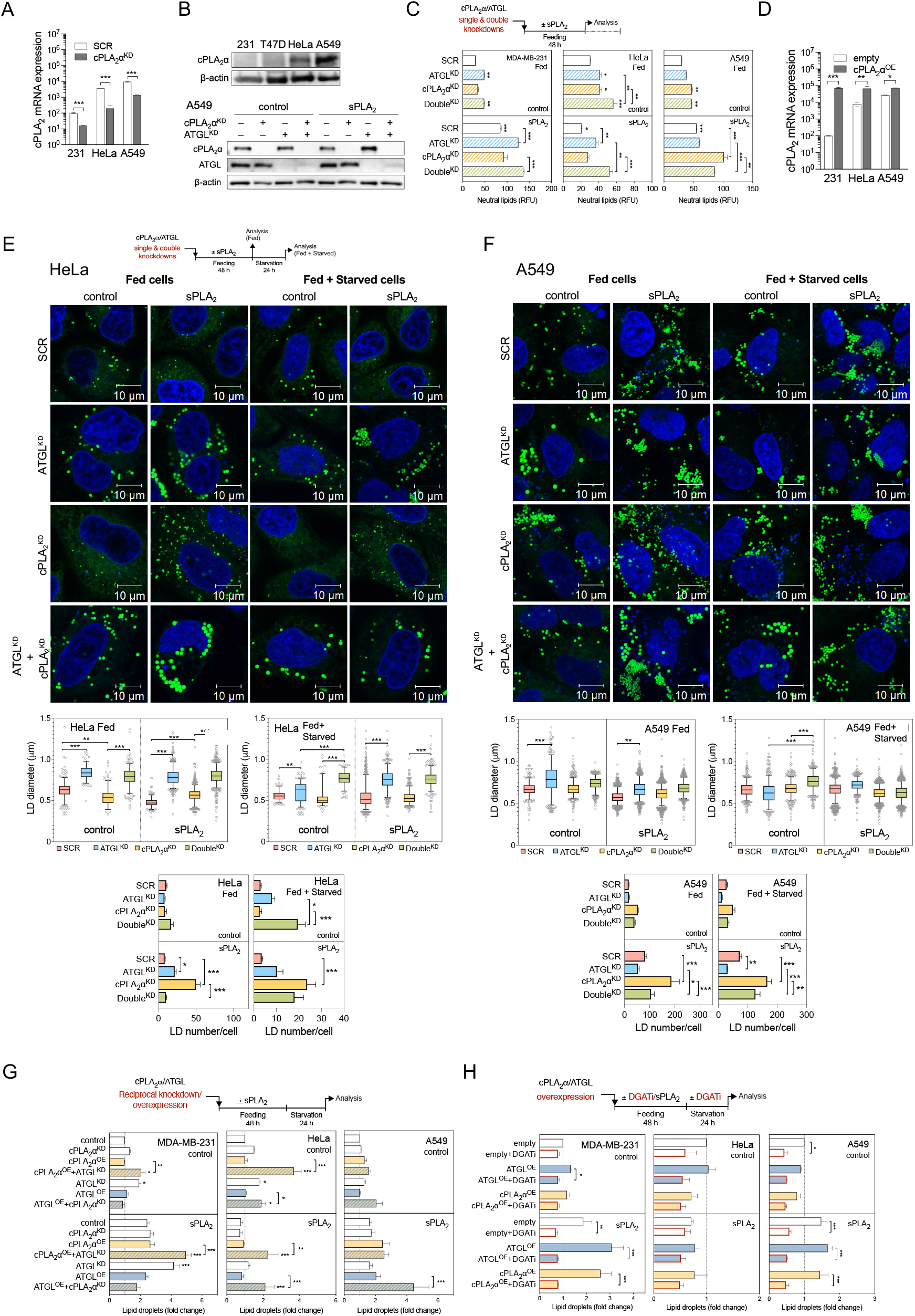
cPLA_2_α cooperates with ATGL in mediating LD-driven lipid mediator production. (**A**) Quantitative PCR analysis of cPLA_2_α gene expression in control (SCR) and cPLA_2_-knockdown (cPLA_2_α^KD^) cells grown for 48 h in complete medium and serum starved for 24 h. (**B**) Representative western blots showing basal levels of cPLA_2_α protein expression in the four cancer cell lines (top) and cPLA_2_α and ATGL protein expression in single and double knockdown A549 cells, without and with hGX sPLA_2_ pre-treatment (bottom). (**C**) Diagram illustrating the experimental conditions used (top), and changes in neutral lipid content in serum-fed cells induced by ATGL (ATGL^KD^) and cPLA_2_α (cPLA_2_α^KD^) single and double (Double^KD^) knockdowns, in comparison with control siRNA-treated cells (SCR), without and with stimulation of LD biogenesis by hGX sPLA_2_ treatment. (**D**) Quantitative PCR analysis of cPLA_2_α gene expression in control (empty) and cPLA_2_α-overexpressing (cPLA_2_α^OE^) cells grown for 48 h in complete medium and serum starved for 24 h. (**E, F**) Diagram illustrating the experimental conditions used, representative live-cell confocal microscopy images and image analysis of LDs in ATGL (ATGL^KD^) and cPLA_2_α (cPLA_2_α^KD^) single and double (Double^KD^) knockdown HeLa and A549 cells, in comparison with non-targeting control siRNA-treated cells (SCR), without and with stimulation of LD biogenesis by hGX sPLA_2_ pre-treatment. LDs were stained with BODIPY 493/503 (green) and nuclei with Hoechst 33342 (blue) and images analysed using ImageJ and the LD Counter Plugin. Box plots are showing changes in LD diameters and LD numbers per cell in serum-fed (Fed) and serum-starved (Starved) cells. Data are geometric means (diameter analysis) or means (number analysis) ±SEM (n >40 cells/sample) of two independent experiments. (**G**) Diagram illustrating the experimental conditions used (top), and neutral lipid levels in cells with reciprocal knockdown/overexpression of cPLA_2_α and ATGL. Cells were reverse transfected with ATGL-targeting (ATGL^KD^) and/or cPLA_2_α-targeting (cPLA_2_α^KD^) siRNAs, then forward transfected with ATGL-encoding (ATGL^OE^) and/or cPLA_2_α-encoding (cPLA_2_α^OE^) plasmids, and/or pre-treated with hGX sPLA_2_. In controls (control), non-targeting siRNA reverse transfections were combined with backbone (‘empty’) vector forward transfections. (**H**) Diagram illustrating the experimental conditions used (top), and DGAT inhibition (DGATi)-induced changes in neutral lipids in serum-starved control cells (empty) and in cells overexpressing ATGL (ATGL^OE^) or cPLA_2_α (cPLA_2_α^OE^), without and with additional stimulation of LD biogenesis by hGX sPLA_2_ pre-treatment. Neutral lipid content was quantified by Nile Red staining and flow cytometry. Data are means ±SEM of two (A, D; C, A549 cells) or at least three independent experiments. *, P <0.05; **, P <0.01; ***, P <0.001 (unpaired t-tests (A, D); two-way ANOVA with Tukey (C, E–G) or Dunnet (H) adjustments; nested one-way ANOVA with Sidak adjustment (E, F)).

**Supplementary Figure 6.**
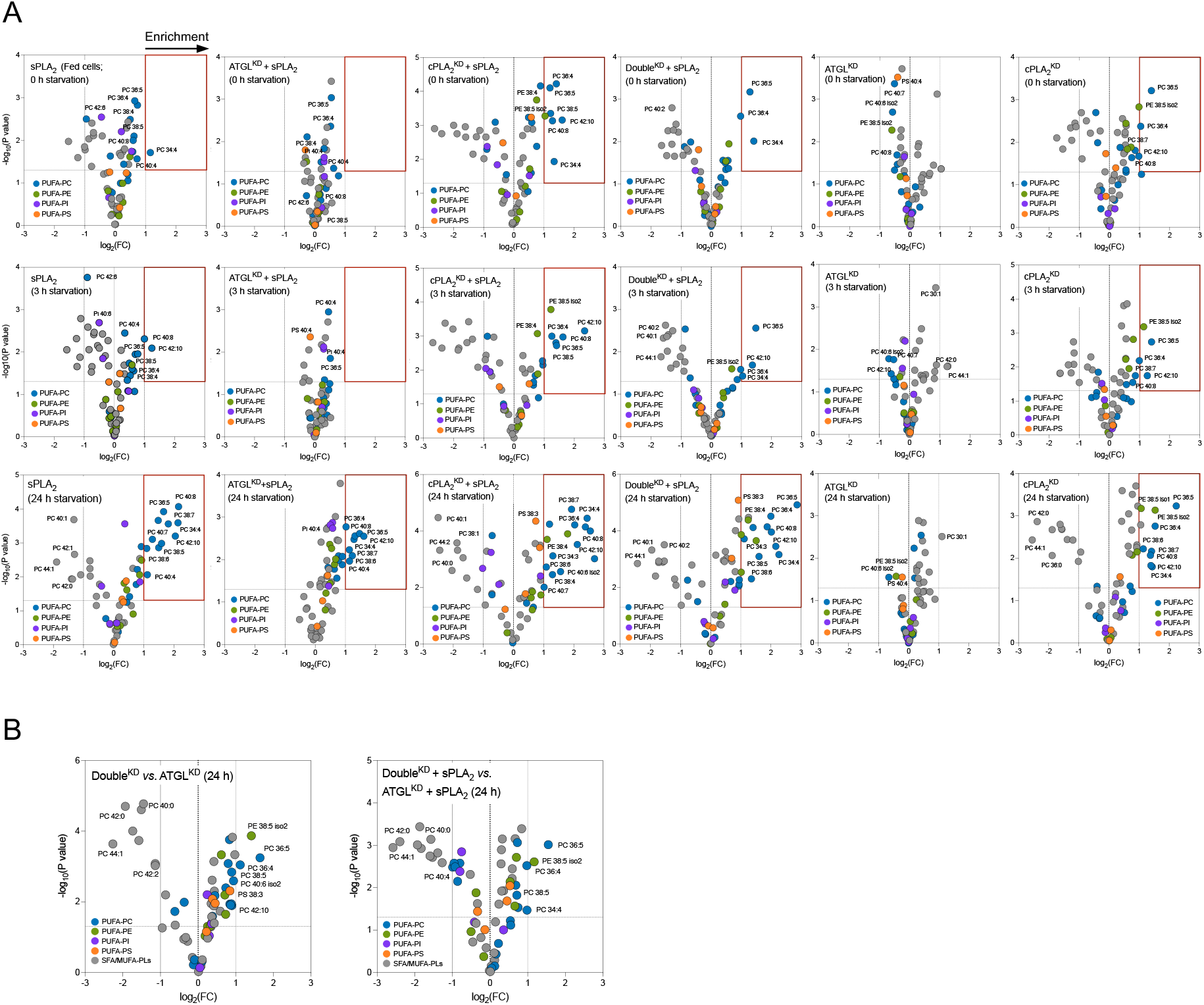
Membrane phospholipid PUFA content is modulated by sPLA_2_, cPLA_2_α and ATGL. (**A, B**) Untargeted lipidomic analysis of phospholipids in MDA-MB-231 cells depleted of ATGL (ATGL^KD^), cPLA_2_α (cPLA_2_α^KD^) or both (Double^KD^), without and with hGX sPLA_2_ pre-treatment, and grown as shown in (B). Volcano plots show significant changes (−log_10_(P value) >1.30) in individual lipids between each treatment condition *versus* control cells (unless otherwise indicated), and were prepared by log_2_ fold-change (FC) data transformation and multiple t-test analysis (n=3 independent experiments). Phospholipids (PLs) containing saturated and mono-unsaturated acyl chains (SFA/MUFA-PLs with 0–2 double bonds) and those containing polyunsaturated FAs (PUFA-PLs with at least 3 double bonds) are colour-coded as indicated. PC, phosphatidylcholine; PE, phosphatidylethanolamine; PI, phosphatidylinositol, PS, phosphatidylserine.

**Supplementary Figure 7.**
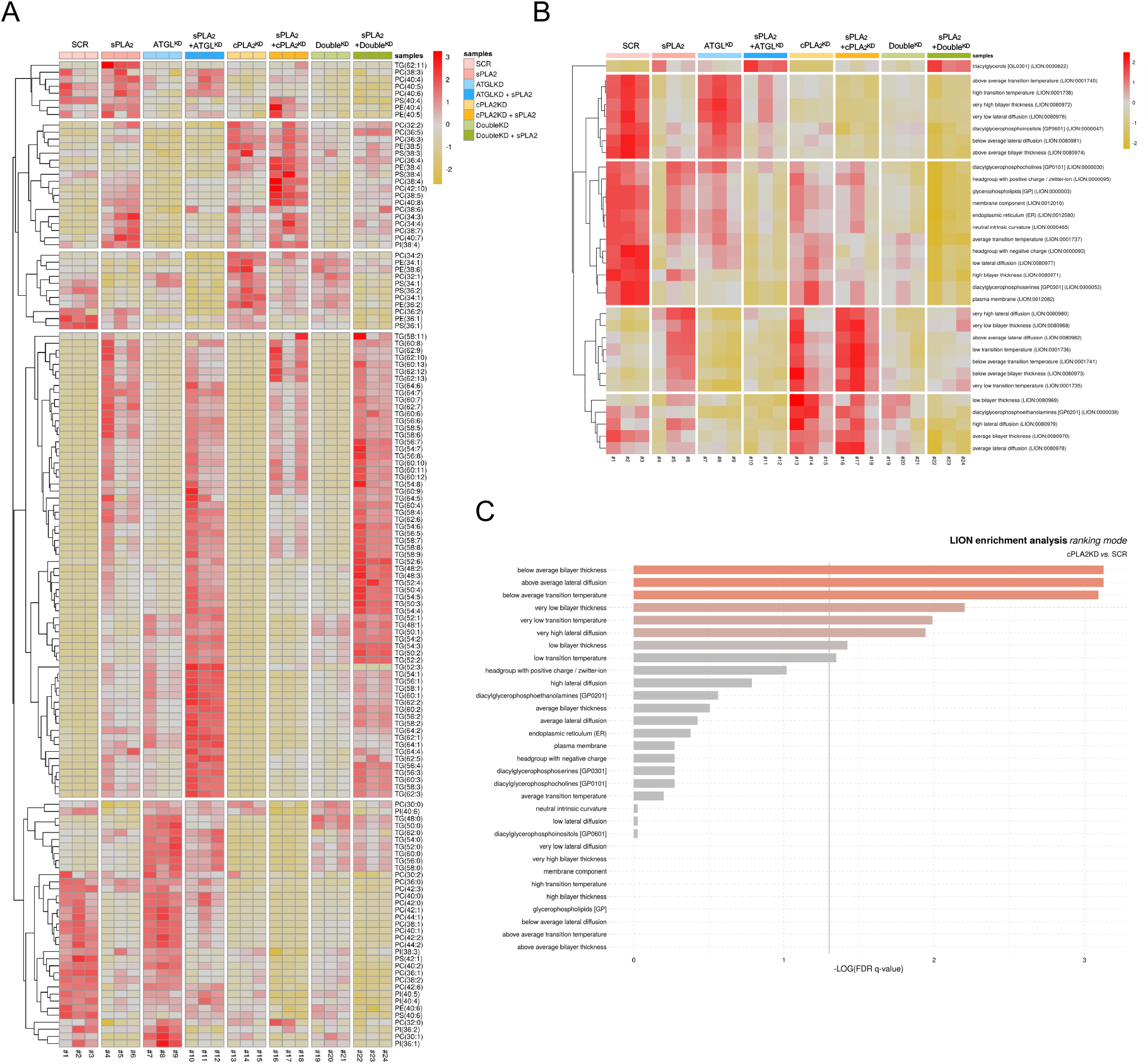
LION-term enrichment analysis for significant changes in membrane properties induced by sPLA_2_, cPLA_2_α and ATGL. (**A**) Heat map of z-score scaled lipid levels of phospholipids and triglycerides in serum-starved MDA-MB-231 cells depleted of ATGL (ATGL^KD^), cPLA_2_α (cPLA_2_α^KD^) or both (Double^KD^), without and with hGX sPLA_2_ pre-treatment under serum-rich conditions. (**B, C**) LION-term enrichment analysis of the full data set (B) and pairwise comparisons of phospholipid data from cPLA_2_α-depleted and control (SCR) cells (C) in ranking mode. The cut-off value of significant enrichments is indicated by the grey line (q <0.05). Data were analysed using three principal components, and lipids were clustered into five groups by hierarchical clustering (A, B). Bar colours are scaled according to the enrichment (−log q-values). FDR, false-discovery rate.

**Supplementary Figure 8.**
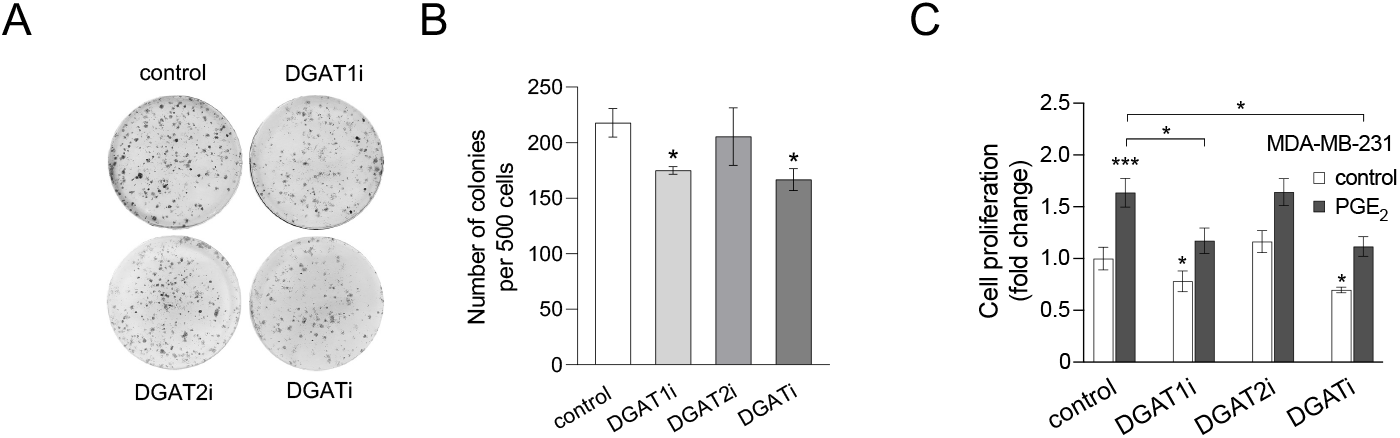
Inhibition of DGAT1 reduces colony formation and PGE_2_-induced proliferation in MDA-MB-231 cells. (**A, B**) Representative images (A) and graph (B) of DGAT inhibition-induced changes in colony formation of MDA-MB-231 cells treated grown in complete medium for nine days and treated with T863 (DGAT1i) and PF-06427878 (DGAT2i), or an equimolar mix of both DGAT inhibitors (DGATi). (**C**) Proliferation of MDA-MB-231 cells treated with 1 μM exogenous PGE_2_ in the absence and presence of T863 (DGAT1i) and PF-06427878 (DGAT2i), or an equimolar mix of both DGAT inhibitors (DGATi). Data are means ±SEM of three independent experiments. *, P <0.05; **, P <0.01; ***, P <0.001 (unpaired t-test (B); two-way ANOVA with Tukey (C) adjustment).

